# Internal model recalibration does not deteriorate with age while motor adaptation does

**DOI:** 10.1101/292250

**Authors:** Koenraad Vandevoorde, Jean-Jacques Orban de Xivry

**Affiliations:** Movement Control and Neuroplasticity Research Group, Department of Movement Sciences, KU Leuven, 3001 Leuven, Belgium; Leuven Brain Institute (LBI), KU Leuven, 3000 Leuven, Belgium

**Author notes:** Corresponding author at: Movement Control and Neuroplasticity Research Group, Department of Movement Sciences, KU Leuven, Tervuursevest 101 bus 1501, B-3001 Leuven, Belgium.; E-mail address (K. Vandevoorde).

**Keywords:** Aging, Motor learning, Motor adaptation, Internal model, Cerebellum

## Abstract

A wide range of motor function declines with aging. Motor adaptation, which occurs when participants learn to reach accurately to a target despite a perturbation, does not deviate from this rule. There are currently three major hypotheses that have been put forward to explain this age-related decline in adaptation: deterioration of internal model recalibration due to age-related cerebellar degeneration, impairment of the cognitive component of motor adaptation, and deficit in the retention of the learned movement. In the present study, we systematically investigated these three hypotheses in a large sample of older women and men. We demonstrate that age-related deficits in motor adaptation are not due to impaired internal model recalibration or impaired retention of motor memory. Rather, we found that the cognitive component was reduced in older people. Therefore, our study suggests the interesting possibility that cerebellar-based mechanisms do not deteriorate with age despite cerebellar degeneration. In contrast, internal model recalibration appears to compensate for deficits in the cognitive component of this type of learning.

## 1. Introduction

The brains of healthy young adults have the ability to quickly adapt motor behaviors to changes in the environment, even if these changes are dramatic (Shadmehr et al., 2010). In contrast, the aging brain is slower to adapt to external perturbations in order to maintain optimal motor performance (Fernández-Ruiz et al., 2000; Seidler, 2006, 2007). Three hypotheses have been put forward to account for this age-related decline in adaptation. Following the widely accepted **internal model hypothesis**, deficits in motor adaptation are due to age-related degeneration of the cerebellum (Seidler, 2006, 2007; Boisgontier and Nougier, 2013; Bernard and Seidler, 2014; Boisgontier, 2015; Hulst et al., 2015; Sugiura, 2016), whose role is crucial for internal model recalibration. Indeed, the cerebellum contains internal models of the body and of the world and makes predictions by transforming motor commands in sensory consequences (Wolpert et al., 1998; Imamizu et al., 2000; Shadmehr and Krakauer, 2008; Shadmehr et al., 2010) and adapting our following movement in order to reduce the sensory prediction error (Shadmehr and Mussa-Ivaldi, 1994; Shadmehr et al., 2010). However, the cerebellum shrinks with aging (Raz et al., 2005), predominantly in the anterior lobe of the cerebellum, involved in motor control (Schmahmann, 2018). A second hypothesis (**strategy hypothesis**) states that recalibration of the internal model is unimpaired by aging (Bock, 2005; Bock and Girgenrath, 2006; King et al., 2013) because some signature of internal model function (Shadmehr and Mussa-Ivaldi, 1994) was identical in elderly adults compared to younger adults (Fernández-Ruiz et al., 2000; Buch et al., 2003; Heuer and Hegele, 2008, 2014; Hegele and Heuer, 2010; Sombric et al., 2017). Moreover, older adults have difficulties acquiring and using explicit strategies to account for the perturbation (Heuer and Hegele, 2008, 2014; Hegele and Heuer, 2010; Huang et al., 2017). Finally, the third hypothesis (**retention hypothesis**) posits that age-related impairments in motor adaptation stem from a deficit in short-term retention (Bock and Schneider, 2002; Trewartha et al., 2014; Malone and Bastian, 2015), which would lead to slower and lesser adaptation (Trewartha et al., 2014).

In the absence of a clear consensus about the mechanisms underlying age-related deficits in motor adaptation, we performed experiments to test each of these hypotheses. To test the internal model hypothesis, single-trial error-based learning (Marko et al., 2012), which drives internal model recalibration, was used to quantify sensitivity to error by presenting visual perturbations of different sizes to participants (Fine and Thoroughman, 2006; Wei and Kording, 2009; Marko et al., 2012; Kasuga et al., 2013; Kim et al., 2018). According to the internal model hypothesis this sensitivity to error should be reduced with aging. Internal model recalibration can also be measured via implicit adaptation in a task-irrelevant clamped feedback task (Morehead et al., 2017). In this task, participants, but not patients with cerebellar degeneration, implicitly adapt their reaching movement to compensate for visual error of constant size (Morehead et al., 2017). According to the internal model hypothesis the implicit adaptation to clamped visual error of constant size should be reduced for older adults as well.

To test the strategy hypothesis, we measured the explicit (i.e. the strategy, which is under cognitive/voluntary control) and the implicit components of adaptation (which remains outside the conscious awareness) (Taylor and Ivry, 2014; Taylor et al., 2014; Haith et al., 2015; McDougle et al., 2015). To measure these two components, we used a cued adaptation task inspired from Morehead et al. in which color cues signaled the presence or absence of the perturbation (Morehead et al., 2015b). The presence or absence of the cues allows participants to switch their strategy on or off (Morehead et al., 2015b). Given that savings, which is the faster relearning of a perturbation, is restricted to the explicit component of adaptation (Morehead et al., 2015a), the strategy hypothesis predicts that both the explicit component of adaptation and savings will be impaired in older adults.

Finally, to investigate the retention hypothesis, we introduced one-minute breaks in the motor adaptation paradigm in order to capture the dynamics of motor adaptation (Sing et al., 2009; Hadjiosif and Smith, 2013). The retention hypothesis suggests that older adults will forget relatively more than young adults during one-minute breaks.

## 2. Materials and methods

### 2.1. Participants

In total 151 healthy adults were recruited and participated after providing written informed consent. In the end 143 of the 151 participants were included in the final analyses. These 143 participants consisted of 72 young adults (between 19 and 36 years old, age: 22.8 ± 2.9 years, mean ± SD; 45 females) and 71 older adults (between 59 and 76 years old, age: 66.8 ± 4.7 years; 35 females). Data of three young participants were excluded before analysis because their data were not saved or because they were not sober at the time of the experiment. Data of five older participants were excluded before analysis because for one the data were not saved properly, one of them did not follow instructions correctly (did not try to adapt during the first learning block) and three of them did not meet the inclusion criteria. The Edinburgh handedness questionnaire (Oldfield, 1971) revealed that all participants were right-handed. All participants were screened with a general health and consumption habits questionnaire. None of them reported a history of neurological disease or were taking psychoactive medication, however 21 older adults reported taking vasoactive medication. In older adults general cognitive functions was assessed using the Mini-Mental State Examination (Folstein et al., 1975). All elderly scored within normal limits (score ≥ 26) (Heuninckx et al., 2008). The protocol was approved by the local ethical committee of KU Leuven, Belgium (project number: S58084). Participants were financially compensated for participation (10 €/h).

### 2.2. Experimental setup

Participants were seated in front of a table and were instructed to make center-out, horizontal reaching movements with their right arm on a digitizing tablet (Intuos pro 4; Wacom). The goal of each reaching movement was to slide through a target with a cursor. The targets and cursor were displayed on a 27 inch, 2560 × 1440 optimal pixel resolution LCD monitor with 144 Hz refresh rate (S2716DG, Dell), vertically mounted in front of the participant. Age-related declines of motor adaptation are observed in both horizontal (Wolpe et al., 2018) and vertical (Heuer and Hegele, 2008) montages of the monitor. Therefore, it is assumed it does not have a big influence on the age effect. Visual feedback was controlled with the Psychophysics toolbox under Matlab. During the reaching movements the participants held a digitizing pen in their right hand as if they were writing. They were instructed to always touch the surface of the tablet with the tip of this pen and to move their right arm and not only their wrist. A wooden cover above the tablet prevented visual feedback from their moving hand. Movement trajectories were recorded at 144 Hz. For this study, four different visuomotor rotation experiments (Figure 1) were designed to investigate the three different hypotheses.

**Figure 1:**
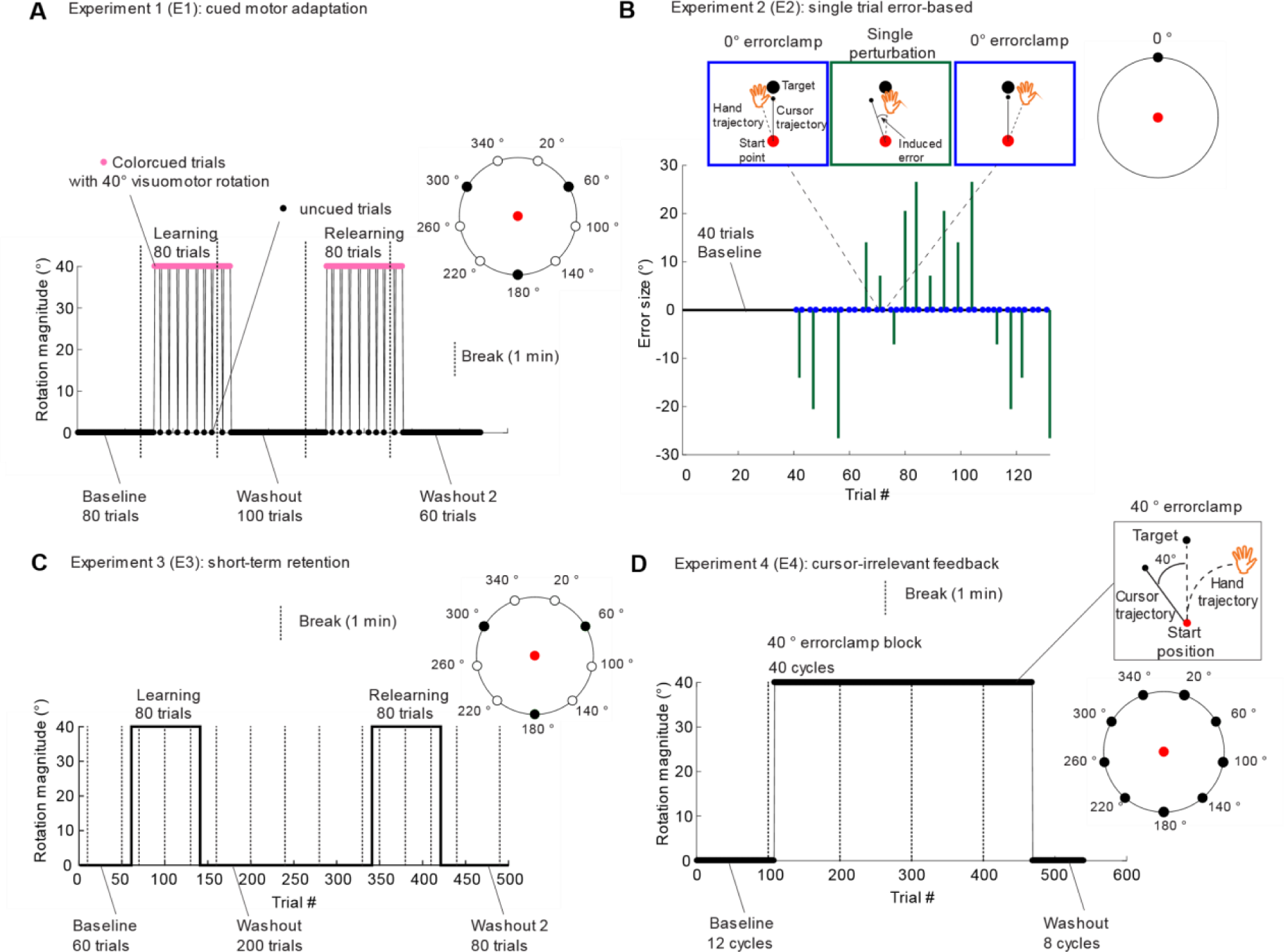
Investigated experiments. **A)** Experiment 1 (E1) to assess the explicit and implicit adaptation level with cue-evoked adaptation. A change in cursor color indicates presence or absence of a perturbation. Nine targets (open black circles) were presented during baseline and washout trial. Three targets (filled black circles) were used during learning blocks. This experiment was replicated with small modifications resulting in two versions of the experiment (E1a and E1b). The number of trials of E1a are indicated in the figure, while E1b consisted of a baseline block of 18 trials, a learning block of 81 trials, a first washout of 99 trials, a relearning block of 81 trials and a second washout of 81 trials. **B)** Experiment 2 (E2) to assess single-trial error-based learning. After 40 baseline trials, participants experienced movement triplets in a random order separated by one or two trials without perturbation. Triplets consisted of a 0 ° error-clamp, one perturbation size and a second 0° error-clamp. In total five possible perturbation sizes were used, both in clockwise or counterclockwise direction (0 °, ± 7.13 °, ± 14.04 °, ± 20.56 ° and ± 26.57 °). A 0 ° error-clamp trial sets the participant’s visual cursor feedback to 0 ° error while the actual movement direction of the hand was not necessarily 0 ° error. The change in actual movement direction of the hand reflects the amount the participant learned from the error experienced in the perturbation trial. The total experiment consists of 10 blocks in which each rotation magnitude is repeated twice (block 1 and 2 are shown). A single target was used during all trials at 0 ° position (filled black circle). **C)** Experiment 3 (E3) to assess retention of motor memory with one minute breaks. Nine targets (open black circles) were presented during baseline and last part of the second washout. Three targets (filled black circles) were used during learning blocks, first washout block and first part of the second washout. **D)** Experiment 4 (E4) to quantify implicit adaptation with task-irrelevant clamped feedback. Nine targets (filled black circles) were used during the baseline, learning and washout blocks.

**Figure 2:**
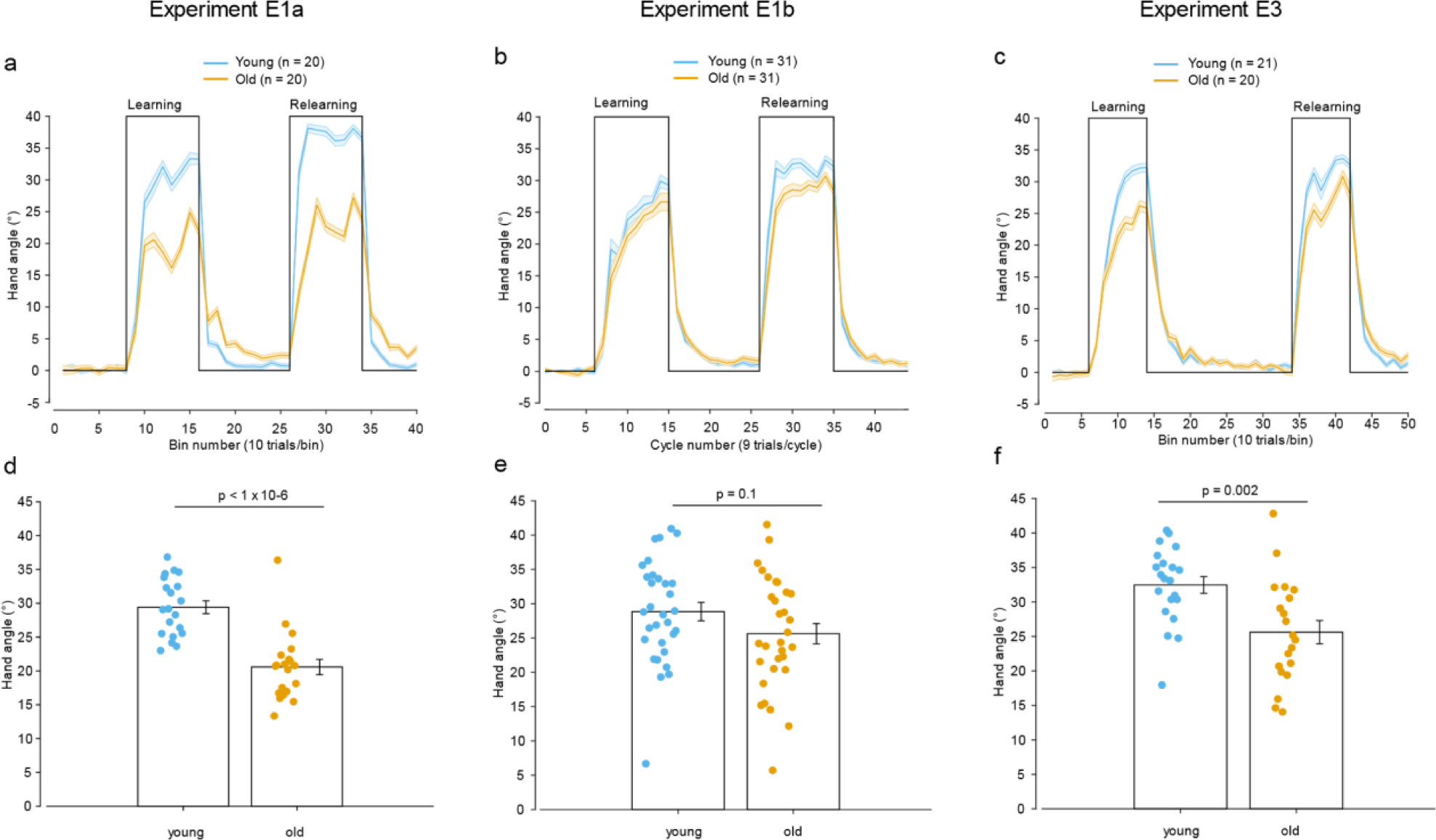
Final adaptation level is decreased in two out of three adaptation experiments for older (in orange) compared to younger adults (in blue). Learning curves for **A)** experiment E1a, **B)** experiment E1b and **C)** experiment E3. Final adaptation level decreased for older compared to younger participants (learning block) for two out of three experiments: **D)** experiment E1a, **E)** experiment E1b (non-significant decrease), **F)** experiment E3.

### 2.3. Assessing implicit and explicit adaptation (Experiment E1; E1a and E1b)

Experiment 1 (E1) was designed to test for both the internal model hypothesis and the strategy hypothesis by assessing implicit and explicit adaptation. It was a visuomotor rotation experiment, adapted from experiment 4 of (Morehead et al., 2015b) with a perturbation magnitude of 40 °. The experiment consisted of a baseline block followed by two pairs of adaptation and washout blocks (Figure 1A). During baseline and washout blocks, we used nine targets spaced 40° apart (from 20° to 340°). During adaptation blocks, three targets were used, spaced 120° (60°,180°,300°) (filled black targets in Figure 1A). About every 80 reaching trials a break of 60 seconds was introduced. The cursor dot remained white the entire baseline and washout blocks. However, during the two adaptation blocks, the cursor became a pink square (i.e. cued trial) instead of a white cursor dot. This cue indicated the presence of a 40° rotation. In each adaptation block, the cursor became again a white cursor dot (i.e. uncued trials) for a few trials, indicating the absence of the perturbation. The instructions were: “First, the cursor will be a white dot, but sometimes the cursor will change to a pink square. At that moment something special will happen but you still have to try to do the same thing, reach to the target with the cursor.” The change in behavior induced by the cue was thus a measure of the explicit component of adaptation as participants could use the cue to switch off any conscious strategies they were applying to counteract the perturbation (Morehead et al., 2015b).

Two versions of experiment E1 were performed: E1a and E1b. In E1a, the first 80 trials were baseline trials, the two adaptation blocks were composed of 80 trials, the first washout block was 100 trials long and the final washout block 60 trials. The length of each block was similar to the length of blocks used in Morehead et al., 2015. Targets were presented purely randomly. There were eight uncued trials in each adaptation block. These eight uncued trials were trials 20, 33, 36, 40, 46, 52, 68 and 72 of the two adaptation blocks of 80 trials. In this version of experiment E1, the presence of an uncued trial should be detected by the participants themselves by carefully observing the cursor shape and color. There was no sound or text to indicate the presence of the uncued trials. Finally, in E1a, the target exploded after hitting it with the cursor, i.e. it became bigger and returned back to normal size together with an explosion sound. No maximum waiting time was imposed between trials.

In E1b, five aspects of the experimental design were modified. First, the targets were presented pseudo-randomly in cycles during baseline and washout with each of the nine targets presented once per cycle. Before the baseline block, participants performed a short dual-task experiment that consisted of a target reaching task and a cognitive reaction time task, results of the dual-task are beyond the scope of the present paper. The adaptation paradigm consisted of a baseline block of two cycles, a learning block, a first washout of 11 cycles, a relearning block and a second washout of nine cycles. In the learning blocks, 9-trial-cycles consisted of three 3-trial-subcycles because only three targets were used. In each subcycle each of the three targets was presented once. Both learning blocks consisted of 9 cycles (or 27 subcycles or 81 trials). Second, we decided to reinforce the awareness of cue switches (signaling a cued trial among uncued ones or an uncued trial among cued ones) with a warning sound played for each cue switch and with a text that indicated the cue switch, displayed for 5 s: ‘Attention! The color of the cursor has changed.’ Moreover, instructions were clearer in experiment E1b: “The trials with a white dot are normal reaching trials like in baseline. The trials with a pink square are special trials. The cursor will often change between a white dot and a pink square.” Third, nine uncued trial were presented per adaptation block (trials 7, 16, 25, 35, 45, 53, 61, 72 and 81). These uncued trials were equally distributed among the three targets (three uncued trials per target). Fourth, no explosions were used when hitting the target. Fifth, a maximum waiting time of 5s was implemented. If participants waited longer, the next trial was initiated. With these extra changes implemented in E1b, we think a better measure of implicit and explicit adaptation could be obtained in E1b compared to E1a for both young and older adults.

### 2.4. Assessing single-trial error-based learning (E2)

Experiment 2 (E2) was another experiment designed for testing the internal model hypothesis. In E2 we assessed error-based learning, therefore we adapted the experiment developed by (Marko et al., 2012) (Figure 1B). Participants experienced first 40 baseline trials with continuous cursor feedback. After these 40 baseline trials, single perturbations could be randomly interspersed throughout the experiment. Perturbations were visual perturbations of different possible angular rotation, i.e. the cursor trajectory is rotated with a given angle with respect to the hand trajectory (Figure 1B). The possible angular rotations were 0 °, ± 7.13 °, ± 14.04 °, ± 20.56 ° and ± 26.57 °. These angles are of similar sizes as in (Marko et al., 2012). Each perturbation angle was experienced 10 times by each participant in both the clockwise and the counterclockwise direction. Before and after each perturbation, error-clamped trials were applied. In error-clamp trials the cursor trajectory was constrained to a straight line from the starting location to the target, regardless of the direction of the reaching movement the participant made (Shmuelof et al., 2012). Cursor distance matched the distance of the hand from the start circle. At the same time, during these error-clamp trials the real reaching movement and hand angles were being registered. Therefore, these error-clamped trials allowed us to measure the reaction to the specific error-sizes as a change in hand angle. In total, this experiment contained 493 reaching trials. The same reaching target was presented throughout the experiment (Figure 1B). This target was positioned 10 cm above the central starting position (at 0 ° direction), away from the body of the participant. About every 80 reaching trials a break of 60 seconds was introduced.

### 2.5. Assessing stability of motor memory (E3)

Experiment 3 (E3) was selected for testing the retention hypothesis by assessing stability of motor memory (Figure 1C). The experiment consisted of 500 trials with a baseline block of 60 trials, two adaptation blocks of 80 trials separated by a washout block of 200 trials and a final washout of 80 trials. Every 30-40 trials a one-minute break was applied (i.e. before trial 10, 50, 70, 100, 130, 160, 200, 240, 280, 330, 350, 380, 410 and 440). These one-minute breaks were used to study stability of motor memory (Hadjiosif and Smith, 2013). They allow us to separate the stable part of the motor memory (remaining motor memory after the break) from the overall part (learning level before the break).

Three targets (60°, 180°, 300°) were used during the adaptation blocks, first washout block and first half of second washout block (Figure 1C). During baseline and during the second half of the second washout block nine targets spaced 40° were used. Similarly as in E1a, the target would explode, accompanied with an explosion sound, after hitting it with the cursor.

### 2.6. Direct measure of implicit adaptation (E4)

Experiment 4 (E4) was our third approach for testing the internal model hypothesis. In addition, the breaks during the learning block of E4 allowed to test the retention hypothesis. E4 was adapted from (Morehead et al., 2017) and aimed at assessing implicit adaptation independently of the explicit component. In this task, the cursor direction of motion was made completely irrelevant by dissociating it from the hand direction of motion (Figure 1D). That is, in these task-irrelevant clamped feedback trials, participants were instructed to ignore the cursor that is always rotated 40 ° with respect to the target direction and to try to move their hand accurately towards the target in the absence of relevant visual feedback of hand position. Targets were presented in cycles of nine trials with each cycle consisting of the nine targets presented randomly (Figure 1D). In total, the experiment consisted of 540 trials or 60 cycles: Baseline consisted of 12 cycles, task-irrelevant clamped feedback trials were presented for 40 cycles and washout consisted of eight cycles. One-minute breaks were given to the participants before trial 100, 200, 300 and 400. In baseline, participants could win points for accuracy. However, during the adaptation block, participants were clearly informed that is was not possible to win extra points, because the cursor could never reach the target during these trials. To keep participants motivated during the adaptation block, the amount of remaining trials was visualized on the monitor.

### 2.7. Organization of the experiments

The presented way of the experiments in this paper is different from the chronological way we conducted the experiment. Experiments were organized in three different paradigms. In total 143 participants took part in one of three paradigms; each paradigm consisted of two experimental sessions with a break of one week between the two sessions (Table 1). The 41 young (age: 22.4 ± 1.8; 25 females) and 40 older adults (age: 66.7 ± 4.9; 17 females) of paradigm 1 and paradigm 2 started with the same experiment on error-based learning (E2) in their first session. The second session, one week after the first one, was different for these participants. In paradigm 1, 20 young (age: 22.2 ± 1.6 years; 10 females) and 20 older adults (age: 66.6 ± 4.9 years; 11 females) performed an experiment for assessing explicit and implicit learning (E1a). In paradigm 2, 21 young (age: 22.5 ± 2.1 years; 14 females) and 20 older adults (age: 66.7 ± 5.0 years; six females) performed an experiment to assess stability of motor memory (E3). In paradigm 3, 31 young (age: 23.3 ± 3.8 years; 20 females) and 31 older adults (age: 67.2 ± 4.4 years; 18 females) started in session 1 with a short visual-spatial working memory (WM) task. After this WM task, experiment 1b (E1b) was performed to assess explicit and implicit adaptation. In session 2, participants started with a neuropsychological test, the Repeatable Battery for the Assessment of Neuropsychological Status (RBANS), which quantified five cognitive measures: language, attention and visuospatial abilities, and immediate and delayed memory (Randolph et al., 1998; Duff et al., 2003). After this test, experiment 4 (E4) was executed to assess implicit learning with task-irrelevant clamped feedback (Morehead et al., 2017).

**Table 1:**
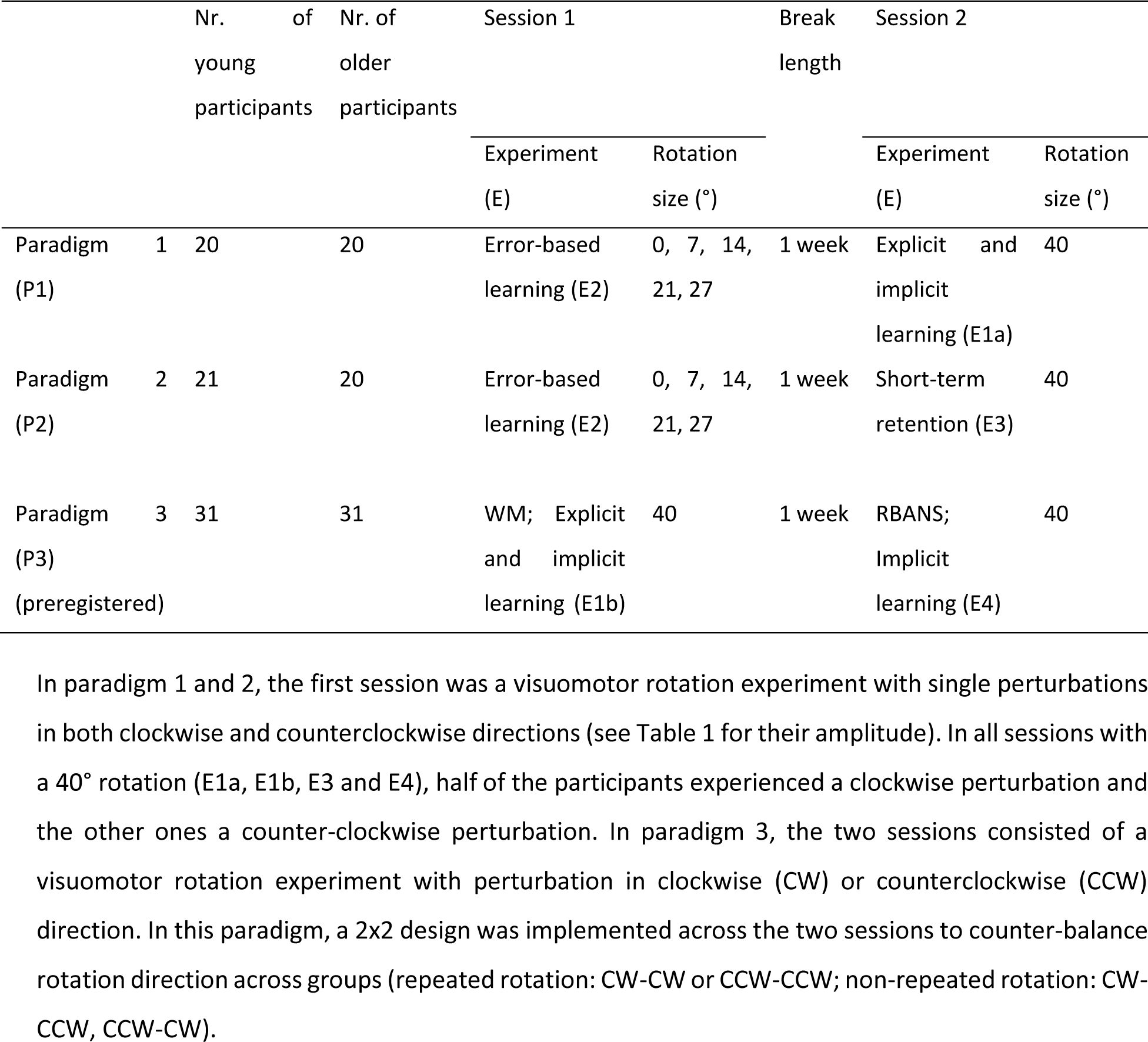
Chronological order of experiments and design of the three different paradigms. Each paradigm (P) consisted of two experimental (E) sessions with a break of one week between the two sessions.

### 2.8. Preregistration

Paradigm 3 was preregistered online: http://aspredicted.org/blind.php?x=3dm6n6. This preregistration included the main hypotheses, the key dependent variables, the amount of participants, the main analyses and secondary analyses investigated with paradigm 3.

The main pre-registered analyses tested for significant differences between the two age groups. Three separate ANOVA’s were applied to analyze the difference in explicit adaptation level (E1b), implicit adaptation level (E4) and the balance of explicit/implicit adaptation level with age group and rotation direction as between-subject factors.

### 2.9. Additional details of the experiments

Each experiment consisted of a series of reaching movements to a single white circular target located 10cm away from the central starting position. For each trial, the participant had to rapidly move his or her right hand to move a white cursor through the target.

The feedback cursor, which represented hand position (when there was no perturbation) was visible until movement amplitude exceeded 10 cm. At this point, a white square marked the position where movement amplitude reached 10 cm, providing visual feedback about the end point accuracy of the reach. The white square had sides of 1.5 mm for E2 and 5 mm for all other experiments. The cursor position froze at the end of each reaching movement and was visible for 1.5 s in paradigm 1 and 2 while it was visible for only 1 s in paradigm 3. All experiments were visuomotor rotation experiments that first started with baseline trials (no perturbation) with normal cursor feedback (i.e. the cursor represents the actual hand position) and continued with perturbation trials where the feedback was either rotated or irrelevant. Before baseline, each participant performed at least nine and maximum 90 familiarization trials to make sure that they understood the instructions and that they performed the task correctly.

There was a different number of targets used depending on the experiment (see below). In paradigms 1 and 2, targets were presented randomly. Therefore, the same target could be repeated several times in consecutive trials. In paradigm 3, targets were presented in pseudo random order. That is, targets were presented in cycles throughout the full experiment and, within each cycle, targets were shown randomly but each target was shown only once per cycle. Therefore, the same target could only be repeated maximum twice in consecutive trials.

In E2, the diameters of the starting point and the target were both 6 mm. The cursor dot had a diameter of 1.5 mm and remained white the entire session. In the other experiments, the diameters of the starting point and the target were both 10 mm and the cursor dot had a diameter of 5 mm.

While returning to the central starting position, the cursor disappeared and only a white circle (i.e. return circle) was visible. The radius of the return circle depended on the position of the pen on the tablet, i.e. the radius of the circle was equal to the radial distance between the position of the hand and the starting point. The center of the return circle was the central starting position. To reduce the time for returning to the starting point, the cursor became visible as soon as the hand was within 3 cm from the central starting position. In paradigm 1 and 2, the return circle was a complete circle. In paradigm 3, an arc was used instead of a complete circle. The reach area was divided in three different zones of 120 °. The arc was in the same 120° zone as where the participant’s (invisible) hand was. Participants had to move their hand in the opposite direction of the arc in order to return to the starting location. In the last 3 cm the cursor did not become visible again. The arc allowed participants to return to the starting position and at the same time prevented the participants from using the visual feedback during the return movement to learn about the perturbation.

Participants were instructed to score as many points as possible by hitting the target with the cursor. When hitting the target, the participant received 50 points. When hitting targets correctly on consecutive trials, 10 bonus points were received for every additional trial with a correct hit (e.g. 60 points were received the second trial after hitting targets correctly on two consecutive trials, 70 points were received the third trial after hitting targets correctly on three consecutive trials). When reaching in close proximity of the target, the participant received 25 points. In E2, the zone for receiving 25 points was an additional 6 mm at both sides of the target. In E2, the zone for receiving 25 points was an additional 5 mm at both sides of the target. The reward for near misses is implemented for keeping participants motivated even when they are not achieving very high accuracies. In E2 and E3 when participants moved too slow, too fast or inaccurate their overall score was reduced with 20 points. Negative overall scores were not possible. In E1 and E4, the overall score could not decrease.

To receive points, participants were required to reach the target between 175 and 375ms after movement onset. If the reaching movement was too slow, a low pitch sound was played and the target color switched from white to blue. If the reaching was too fast, a high pitch sound was played and the target color switched from white to red. The cumulative score of all previous trials was displayed throughout the experiment.

The experimenter (KV) was present during the entire experiment to motivate the participants to achieve the highest possible score and to make sure that the participant performed the task correctly. The experimenter regularly reported that the participant was performing well, even when the score was below average.

At the end of the feedback period, the participant had to move the tip of the pen back to center of the tablet and wait there between 350ms and 850ms (in steps of 50ms) in order to start the next trial. In paradigm 1 and 2, no maximum waiting time existed between the different reaching movements. Therefore, participants could wait, in theory, as long as they wanted to before initiating their reach to the target. In paradigm 3, a maximum waiting time (5 s) was implemented. If participants waited too long, the next trial was initiated.

### 2.10. Cognitive assessment

#### 2.10.1. Repeatable battery for the assessment of neuropsychological status (RBANS)

Neurocognitive status is quantified with RBANS (Randolph et al., 1998). Index scores were obtained for five domains: Immediate memory, visuospatial/constructional abilities, language, attention and delayed memory. Twelve tests are used to quantify these five domains: List learning and story memory for immediate memory, figure copy and line orientation for visuospatial, picture naming and semantic fluency for language, digit span and coding for attention and list recall, list recognition, story recall and figure recall for delayed memory. In list learning, participants have to immediately recall a list of ten words over four learning trials. In story memory, participants have to recall a story twice with 12 main items. In figure copy, they have to draw a geometric figure with 10 parts with the right accuracy and placement. Line orientation consists of 10 trials in which participants have to match two lines with a specific orientation to an array of 13 lines. During picture naming, they have to name 10 different items. In semantic fluency, participants are required to give as many items as possible for a semantic category. Digit span probes the amount of numbers a participant can keep in working memory with the amount increasing from two until nine numbers. The coding task requires the participant to transform a code into numbers as fast as possible. Finally, list recall, list recognition, story recall and figure recall tests all test performance on the list learning, story memory and figure drawing after a period of delay (Randolph et al., 1998). The results of these cognitive tests allow to determine whether correlations exist between individuals’ adaptation level and neurocognitive status. These correlations were purely exploratory.

RBANS consists of 12 tests to quantify five cognitive measures. In normative populations, the raw test scores decline with aging. Therefore, the raw test scores are first converted to index scores for six age-groups. Each age-group includes a span of 10 years, with the exception of the first age-group, which spans 20 years from age 20y until 39y. The obtained index-scores allow comparison of scores across individuals independently of their age. The sum of all index-scores gives the sum-index, which is an overall measure of cognitive performance (Randolph et al., 1998). RBANS index scores could not be used for comparison between the two age groups or for correlation analysis because of the age-normalization, raw RBANS test scores were used instead.

#### 2.10.2. Visuospatial working memory task

A computer-based task was used to quantify visuospatial working memory capacity (WMC). Sixteen white squares (1.9 cm × 1.9 cm) were presented in a circular array (11.2 cm diameter). Three, four, five or six red circles (0.8 cm diameter) were visualized for two seconds in the 16 white squares with each red circle presented randomly in one of the 16 squares. Participants were asked to remember the positions of the presented red circles. After these two seconds, participants fixated on a white cross (0.6 cm × 0.6 cm) for three seconds. Afterwards they were asked to make a button press within three seconds to indicate whether a probed location corresponded to a position that contained a red circle before (McNab and Klingberg, 2008; Christou et al., 2016). After the three seconds for responding, they had to fixate on a small blue cross (0.2 cm × 0.2 cm) for one second. In total, one trial had a fixed time duration of nine seconds. First participants could practice the working memory task with eight trials. After the practice session, each participant had to complete 40 trials. The 40 trials contained three, four, five or six red circles (10 trials/condition) with all conditions randomly mixed.

### 2.11. Data analysis

Analyses of paradigm 1 and 2 were performed without preregistration, while analyses of paradigm 3 were preregistered. All data and analysis scripts can be found on Open Science Framework (https://osf.io/vncce/).

#### 2.11.1. General analysis

All analyses and statistical calculations were performed in MATLAB 2017b (The MathWorks). For each reaching movement, the hand angle (relative to target angle) was calculated from the first data point exceeding 4 cm distance from the middle of the starting point. The time for reaching 4 cm was on average 172 ms in E1a, 144 ms in E1b, 167 ms in E2, 173 ms in E3, 134 ms in E4. The hand angle was the primary dependent variable in all of the experiments. The angular error is the angle the cursor deviated from the target. Angular errors above 60 ° were due to inattentive reaches to previous target directions and were assumed to be outliers. These outliers were removed before processing the data. The statistically significant threshold was set at p<0.05 for the ANOVA’s. We reported effect sizes (partial eta squared: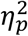) as well as F and p-values.

#### 2.11.2. Analysis 1: Final adaptation level

To assess the final adaptation level of each participant, we averaged the hand angles of the 18 trials of each learning block. These hand angles were first corrected with the average hand angles of the last 18 baseline trials before each learning block. Statistical comparison was performed with three separate 2-way ANOVA’s, one for each experiment. The between-participant factors were the age group (young or old) and the rotation direction (clockwise or counterclockwise). Here, we only report the differences for the learning blocks, the differences in relearning are reported together with the results of explicit adaptation and savings.

#### 2.11.3. Analysis 2: Overall adaptation learning rate

An exponential function (Eq. 1) was fit to the angular errors of all the learning blocks of experiments E1 and E3 (Figure 1).

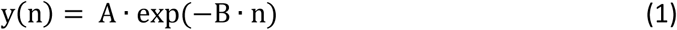

where n is the trial number in the specific learning block and A and B are two constants. The B-parameter is a quantification of the overall adaptation rate. We expected an increased value of learning rate for young compared to older subjects. The distribution of this parameter under the null hypothesis was obtained by computing all the possible values under resampling (N = 10.000) with random reassignment of the participants in two subgroups (without replacement). The p-value was defined as the portion of the resampled distribution that was more extreme than the observed statistic (Hesterberg et al., 2003). Mean and standard deviation of learning rate were obtained after 10.000 bootstraps. This analysis was preregistered as a secondary analysis for E1b.

#### 2.11.4. Analysis 3: Implicit adaptation with cued motor adaptation (E1a and E1b)

The first adaptation block was corrected for baseline errors by subtracting the average error of the last 18 trials of baseline. The second adaptation block was corrected by subtracting the average error of the last 18 trials of washout. We analyzed the data in all the uncued trials that were preceded by a cued trial (eight uncued trials for E1a and nine uncued trials for E1b per learning block). The amount of implicit learning was calculated per learning block as the average of the uncued trials (Morehead et al., 2015b). Two separate 2-way ANOVA’s were used, one for the first and one for the second learning block, with the between-subject factors, age and rotation direction, and with the implicit adaptation as dependent variable.

#### 2.11.5. Analysis 4: Error-based learning (E2)

Hand angles during the error-clamps immediately before and after a trial with an induced error were compared (Figure 1B). Learning was quantified for four different perturbation sizes (7.13 °, 14.04 °, 20.56 ° and 26.57°) and two directions (clockwise and counterclockwise) separately. Learning for 0° perturbation was implemented as a control measure. A repeated measures 3-way ANOVA was used with the within-subject factors, perturbation size and the rotation direction (CW and CCW), and the between-subject factor, age (young and old). The reaction to 0° error size was analyzed with a separate unpaired 2-sided t-test to make the comparison between young and old.

In the error-based learning experiment (E2), single errors were introduced (induced errors) to the participants (Figure 1B). However, on such perturbation trials participants could still make errors themselves. This creates experienced errors which are slightly different from induced errors. Therefore, the previous analysis was repeated with the actual errors instead of the induced errors. To see more details about the error correction, we chose to double the amount of bins compared to the original amount of induced error sizes. This resulted in eight bin sizes from 0° until 26.57°. Thereafter, a repeated measures 2-way ANOVA was used with the within-subject factor, the experienced error size bin (0° to 3.57°, 3.57° to 7.13 °, 7.13 ° to 10.70°, 10.70° to 14.04 °, 14.04 to 17.61°, 17.61° to 20.56 °, 20.56 ° to 24.13 °, 24.13 ° to 26.57 °) and the between-subject factor, age (young and old).

#### 2.11.6. Analysis 5: Implicit adaptation with task-irrelevant clamped feedback (E4)

The amount of implicit learning was the average of the last 10 cycles of the learning block. The adaptation block was first corrected for baseline errors by subtracting the average error of the two last baseline cycles (18 trials). A 3-way ANOVA was used for analysis with the between-subject factors, age, rotation and congruency (preregistered as a primary analysis). We could verify whether congruent perturbation directions in two sessions influenced the adaptation level. Congruency defines whether the perturbations in the two sessions of paradigm 3 (E1b and E4) were in the same direction or not.

Finally, the rate of implicit adaptation was compared between young and older adults. Therefore, we fitted an exponential function (Eq. 1) to the baseline-corrected adaptation block (360 trials) and performed 10.000 resamplings as described in analysis 2 (preregistered as secondary analysis). Mean and standard deviation of learning rate were obtained after 10.000 bootstraps.

Additionally, Pearson correlations between implicit adaptation from the cued motor adaptation experiment and the implicit component from the task-irrelevant clamped feedback were calculated (Figure 3-1) (preregistered as secondary analysis for E1b and E4). Correlations were determined for the combination of the same and opposite rotation directions, only the same rotation directions and only opposite rotation directions across two sessions. This resulted in six correlations which were corrected for multiple testing with false discovery rate (FDR) (Benjamini and Hochberg, 1995).

In addition to correlation analysis, a robust linear regression (robustfit in Matlab) was performed in order to verify that correlations were not influenced by between group differences in these variables. Task-irrelevant implicit adaptation (Y) was estimated using a linear combination of implicit adaptation from E1b (X), a binary age vector (G) and the interaction of X and G in the regression equation with intercept A and regression coefficients (B,C,D):

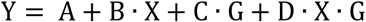

Standardized beta coefficients could be obtained instead of regression coefficients when first converting variables X and Y to z-scores and afterwards applying linear regression.

#### 2.11.7. Analysis 6: Explicit adaptation (E1a and E1b)

Baseline subtraction of the two learning blocks was the same as in analysis 5. The amount of explicit learning was calculated by subtracting hand direction in the uncued trials (see analysis 3) from the cued trials immediately preceding those (Morehead et al., 2015b). Two separate 2-way ANOVA’s were used to analyze the first and second learning block with the explicit adaptation level as the dependent variable and with the between-subject factors, age and rotation. The 2-way ANOVA to analyze the first learning block of experiment E1b was preregistered as a primary analysis. To analyze savings of explicit adaptation, a 3-way ANOVA was used with the amount of explicit learning as the dependent variable, the within-subject factor, the learning block, and the between-subject factors, age and rotation.

#### 2.11.8. Analysis 7: Cue-evoked savings (E1a and E1b)

In addition, cue-evoked savings was calculated as the difference in baseline-subtracted hand angles in the first trial of the relearning block compared to the first trial of the learning block (Morehead et al., 2015b). A 2-way ANOVA was used for analysis with the between-subject factors, age and rotation direction, and the dependent variable, cue-evoked savings.

#### 2.11.9. Analysis 8: Balance explicit/implicit adaptation (E1b and E4)

The balance of explicit/implicit adaptation is calculated as the amount of explicit adaptation as calculated in analysis 6 divided by the amount of implicit adaptation as specified in analysis 5. A 4-way ANOVA was used with the explicit/implicit adaptation balance as the dependent variable, with age, rotation and congruency as between-subject factors and with learning block as within-subject factor. This balance calculation is highly impacted by the variability of implicit adaptation in the denominator, which induces high variability. However, it was specified in the preregistration as a primary analysis for E1b and therefore we added the results in Figure 5-1.

#### 2.11.10. Analysis 9: Correlation analysis between adaptation components (E1a and E1b)

We calculated the correlations between different measures of the adaptation process (overall, explicit and implicit adaptation) for participants from experiments E1a and E1b with Pearson correlation coefficients. Because these three measures of adaptation are not fully independent when computed on the same learning period, we correlated measures from the learning and relearning periods. Robust outlier removal was performed according to Pernet et al (Pernet et al., 2013). The correlation analysis between the explicit and implicit adaptation was preregistered as a secondary analysis for E1b.

In addition to correlation analysis, a robust linear regression (robustfit in Matlab) was performed in order to partial out the effect of age group on the correlations as explained in Analysis 5.

#### 2.11.11. Analysis 10: Working memory capacity (WMC)

The computer-based working memory task allows to determine WMC with the K-value, estimating the number of items that can be stored in WM (Vogel et al., 2005), calculated as K = S(H-F) using three to six items. This is similar to the original experiment (Vogel et al., 2005) but differs from what previous adaptation studies (Christou et al., 2016) have used where the K-value (i.e. K56) was obtained from the trials with five and six items only. We chose to measure WMC with all items because it was not possible to replicate the correlation between WMC and explicit adaptation for young participants as mentioned in (Christou et al., 2016) with the WMC measure with only five and six items. This correlation with WMC measured with all items and overall adaptation was also significant in the study of Christou (personal communication from Dr. Galea). Two separate 1-way ANOVA’s with between-subject factor, age, were executed to assess age differences for the K and K56 values (preregistered as a secondary analysis for E1b and E4).

#### 2.11.12. Analysis 11: Neuropsychological status

For each of the 12 individual cognitive RBANS tests each participant receives a score. A repeated measures 2-way ANOVA was executed with between-subject factor, age, and 12 repeated measures, one for each RBANS raw test score. This analysis was preregistered as a secondary analysis for E1b and E4.

#### 2.11.13. Analysis 12: Correlation analysis with cognitive measures

Spearman correlation analyses were performed between individual’s explicit adaptation levels (E1b) and the K-value, and between individual’s explicit adaptation levels (E1b) and the RBANS cognitive raw test scores. Outlier removal proceeded according to analysis 9. This analysis was preregistered as a secondary analysis for E1b and E4.

The two working memory capacities (K and K56) were related to two explicit adaptation components which resulted in four correlations. In the correlation analysis, we corrected for multiple comparisons by calculating the adjusted p-values with FDR.

Five cognitive measures (figure copy, digit span, coding total, figure recall and line orientation) were related to two explicit adaptation components which resulted in 10 correlations. In the correlation analysis, we corrected for multiple testing by calculating the adjusted p-values with FDR. In preregistration, the RBANS correlation analysis also mentioned correlations with immediate and delayed memory index scores. However, these correlations are only given in supplementary Figure 5-1 because it made no sense to use age-normalized index scores.

In addition to correlation analysis, a robust linear regression (robustfit in Matlab) was performed in order to partial out the effect of age group on the correlations as explained in Analysis 5.

#### 2.11.14. Analysis 13: Short-term relative retention of motor memory

Retention of motor adaptation was quantified with a visuomotor rotation experiment (E3) with three breaks of one minute in each learning block. The stable components of adaptation (*x*_*stable*_) are the hand angles after the breaks in the two learning blocks (Trials 11, 41 and 71 after the onset of perturbation). To estimate the level of overall adaptation before the breaks, we averaged the level of adaptation of one bin (10 trials) before the breaks (*x*_*overall*_, computed for trials 1-10, 31-40 and 61-70 after the onset of perturbation). The stable components were expressed as percentages of the overall components of motor adaptation. This percentage indicates the relative retention of motor memory with the formula:

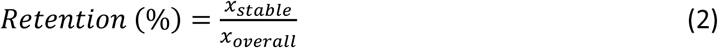

As three breaks were present in each adaptation block, six values for relative retention were calculated for each participant. A repeated measures 4-way ANOVA was used with between-subject factors, age and rotation direction, and with within-subject factors, the two learning blocks and the three breaks in each learning block. The dependent variable was the retention of motor adaptation.

In addition, the relative retention (%) of motor adaptation was quantified for the task-irrelevant clamped feedback experiment (E4) with formula (2). Three breaks were applied in the adaptation block of experiment E4 (trial 200, 300 and 400). The overall (*x*_*overall*_) and stable (*x*_*stable*_) components of adaptation were calculated for each of these three breaks by averaging the hand angles of one complete cycle of nine targets, respectively before and after the breaks. Given that the trials immediately before and after the breaks were not directed to the same target, we selected a complete cycle before and after the break. A repeated measures 3-way ANOVA was used with between-subject factors, age and rotation direction, and with within-subject factor, the break number in the adaptation block. The dependent variable was the retention of motor adaptation.

#### 2.11.15. Analysis 14: Relative change in reaction time and other reaching variables

To assess the relative change in reaction time during motor adaptation, we averaged the reaction time of the last 18 trials of each learning block. These were normalized for each participant by subtraction of the average reaction times of the last 18 baseline trials before each learning block. Statistical comparison was performed with seven separate 1-way ANOVA’s, one for each experiment and one for each learning block, the between-subject factor was the age group.

Nine unpaired 2-sided t-tests were executed for each experiment to compare young and older participants for eight dependent variables of the experiments. The dependent variables were trial time duration (in s), reaching time (in s), maximum displacement, surface displacement, number of too fast trials, number of too slow trials, total score, inter-trial interval (in s) and total duration of experiment (in s). Trial time duration is the time from the start of the reaching movement to the end of it. Reaching time was calculated as the time point when participants exceeded 4 cm distance. The inter-trial interval is the time between the end of the reaching and the start of the next reaching movement. Maximum displacement and surface displacement are both indicating how curved the reaching movements were. To obtain these variables for one reaching trial, a straight line was drawn between the start and end point of the reaching movement. The maximum displacement is the maximum distance from the reaching movement to the straight line and the surface displacement is the surface between the reaching movement and the straight line (preregistered as secondary analysis for E1b and E4). These variables were selected because they give a good overview of the parameters that are affected by age.

## 3. Results

### 3.1. Aging affects final adaptation level and the adaptation rate

Young (N = 72, age = 22.75 y ± 2.85 SD) and older (N = 71, age = 66.88 y ± 4.65 SD) participants performed reaching movements under visuomotor rotation. In three experiments (Figure 1, E1a, E1b and E3), we investigated the difference in adaptation between young and old participants which typically declines with aging (Seidler, 2006, 2007; Heuer and Hegele, 2008). In these three experiments, we confirmed that older adults adapted less than younger adults (Figure2A-C) although the extent of this decline was quite variable across experiments. To quantify the effect of age on motor adaptation, we measured its extent by looking at hand angles over the last 18 trials of the first learning block (analysis 1). The effect of age on the second (relearning) block will be evaluated in a later section (see below).

In experiments E1a and E3, we observed that the final adaptation level was significantly lower for older adults (analysis 1) (E1a (Figure 2A): F(1,37)=22.9, p < 1×10^−6^, 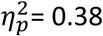 E3 (Figure 2C): F(1,38)=8.7, p = 0.005, 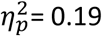) but not in experiment E1b (E1b (Figure 2B): F(1,58)= 2.49, p =0.1, 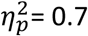).

Beyond the final adaptation level, some studies also suggest that the rate of motor adaptation is decreased for older adults compared to younger ones (Fernández-Ruiz et al., 2000; Anguera et al., 2011). To compare the learning rates, we fitted exponential functions to the angular errors during learning blocks and performed a permutation test (n = 10.000) for each comparison (analysis 2). In line with these previous studies and with the observation of the affected final adaptation level, we found an affected learning rate for older adults compared to younger ones in two motor adaptation experiments (analysis 2)[E1a: p < 0.0001; E3: p = 0.0009](E1a; mean ± std: young: 0.033 ± 0.006, old: 0.008 ± 0.001; E3; mean ± std: young: 0.031 ± 0.004, old: 0.016 ± 0.002). Also in line with the result of the final adaptation level, the learning rate was not affected for older adults in experiment E1b (analysis 2) [E1b: p = 0.5](E1b; mean ± std: young: 0.018 ± 0.003, old: 0.015 ± 0.002). Overall, aging affected final adaptation and learning rate.

### 3.2. Internal model recalibration does not deteriorate with age

While the overall adaptation was sometimes decreased in elderly people, this does not reveal which component of adaptation is affected. Therefore, we tested whether this deterioration of performance was due to their inability to recalibrate their internal model (implicit adaptation). Such recalibration of internal models can be measured via its measure during motor adaptation ((Morehead et al., 2015b); Figure 3A-D), via single-trial learning ((Marko et al., 2012); Figure 3E-F) or via the drift elicited by sensory prediction errors ((Morehead et al., 2017); Figure 3G-H). The impact of aging on each of these measures is reported below.

**Figure 3:**
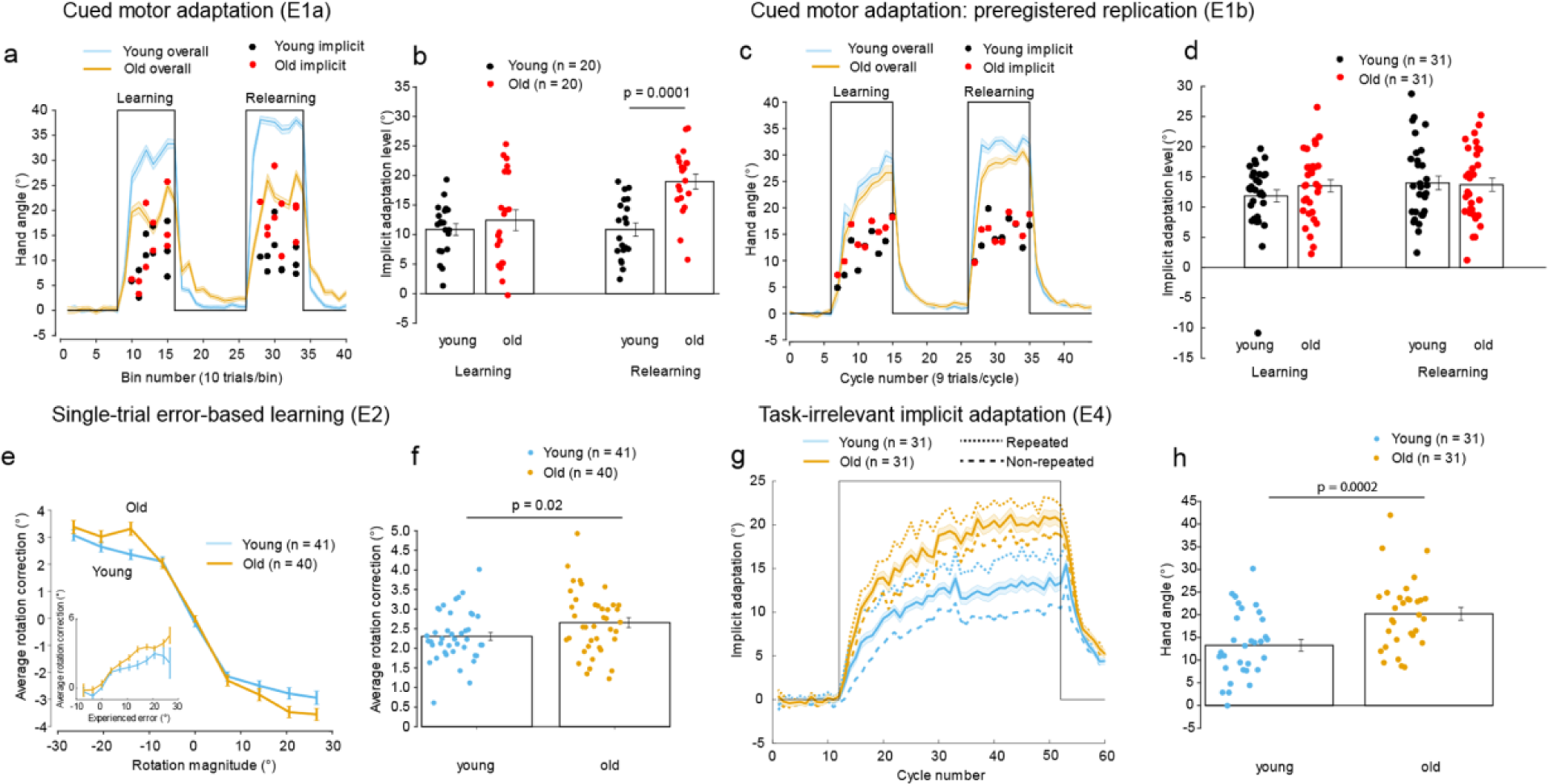
Internal model recalibration does not deteriorate with age. **Results of cued motor adaptation (E1): A)** Decreased overall cue-evoked adaptation for older adults, as measured with E1a. During uncued trials implicit adaptation was measured, visualized with red (older adults) and black (younger adults) dots. **B)** Same implicit data as fig 3A. Visualization of individual data points for implicit adaptation. **C)** Overall cue-evoked adaptation for older adults, as measured with E1b. During uncued trials implicit adaptation was measured, visualized with red (older adults) and black (younger adults) dots. **D)** Same implicit data as fig 3C. Visualization of individual data points for implicit adaptation. **Results of single-trial error-based learning (E2): E)** On average older adults react more to specific single induced error sizes. These data were obtained with E2, described in fig 1B. The inset shows that older adults react more to experienced errors as well. **F)** Same data as fig 3E, but now the Individual data points for reaction to error are shown after averaging reaction to error for error sizes (7.13 °, 14.04°, 20.56° and 26.57°). **Results of task-irrelevant clamped feedback experiment (E4): G)** On average older adults adapt more implicitly, as measured with E4, described in fig 1D. Implicit adaptation for repeated perturbation is higher than for non-repeated perturbation both for young and older adults. **H)** Same data as fig 3G, but now individual implicit adaptation levels are shown. See also Figure 3-1.

Implicit adaptation was quantified in experiment 1, a cued motor adaptation experiment (Figure 1A; E1), in which color cues allowed participants to voluntarily switch their explicit aiming on or off. In the first learning block of E1a, we observed no difference in implicit adaptation across age (analysis 3) (E1a (Figure 3A-B): F(1,36) = 1.9, p = 0.2, 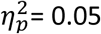; E1b (Figure 3C-D): F(1,58) = 1.65, p = 0.2, 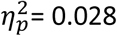). In the second learning block, we observed a significant increase in implicit adaptation level for older adults in the first experiment (analysis 4) (E1a (Figure 3A-B): F(1,36) = 19, p = 0.0001, 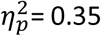) but not in the second experiment (E1b (Figure 3C-D): F(1,58)= 0.02, p = 0.9, 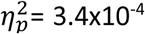). For both learning blocks, no interaction between age and rotation direction was observed for the first and second experiment. Even though the statistical results differ across experiments, both indicate that the implicit adaptation of elderly participants was at least as good as that of younger participants.

Single-trial learning was quantified in experiment 2 (E2) by looking at how participants changed their behavior after a single perturbation trial (Figure 3E-F). To do so, we compared the behavior in trials before and after the participants experienced a visuomotor rotation of one of five possible angles on a single trial. As illustrated in Figure 3E and 3F, change in movement direction was increased with increasing perturbation sizes (analysis 4) [main effect of perturbation size: F(3,237) = 18.8, p < 10^−10^] and for older compared to younger adults [F(1,79) = 5.45, p = 0.02, 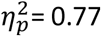]. In addition, the rotation correction for the control perturbation size of 0° was not different for young and old [t(79) = −0.98, p = 0.33] which shows that the bias of reaching without a perturbation was similar across groups. Rotation direction had no effect on reaction to single error sizes [F(1,79) = 0.15, p = 0.7] and no significant interactions between age and the other factors were present [age x error size: F(3,237) = 1.5, p = 0.2; age x rotation: F(1,79) = 0.02, p = 0.9; age x error size x rotation: F(3,237) = 1.2, p = 0.3]. When the change in hand angle was analyzed using the experienced error sizes instead of the induced error sizes, the age-effect of increased reaction to error is still present (analysis 4; inset in Figure 3E) [F(1,79)= 294.3, p <10^−27^, 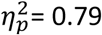]. In summary, these results show that older adults react more to errors than young adults.

During the task-irrelevant clamped feedback experiment (Figure 1D; E4) participants experienced a constant visual error that was irrespective of their movement, which induced a drift in the direction opposite to the cursor motion (Figure 3G). This change in movement direction was larger for older compared to younger adults (Figure 3H; F(1,54) = 15.6, p = 2 × 10^−4^, 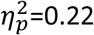) (analysis 5). The direction of the perturbation did influence the adaptation as change in movement direction was higher for a clockwise than for a counter clockwise 40 ° perturbation [F(1,54) = 9.8, p = 3 × 10^−3^]. In addition, we analyzed the influence of the retrograde interference between the two sessions (E1b and E4, separated by one week) on our results. Participants who received the same perturbation direction (i.e. repeated direction) in both sessions reached a higher level of implicit adaptation in E4 than those who received two opposite perturbations (i.e. non-repeated) (analysis 5, Figure 3G) (main effect of congruency, F(1,54) = 6.8, p = 0.01). However, this carry-over effect does not affect our conclusion as the effect was similar for congruent and incongruent condition (see Figure 3G), Indeed, we found no significant interactions between age and congruency [age x rotation: F(1,54) = 0.8, p = 0.4; age x congruency: F(1,54) = 0.1, p = 0.8; age x rotation x congruency: F(1,54) = 0.2, p = 0.7]. The implicit component from the cued adaptation experiment (E1b) was correlated with the implicit adaptation in cursor-irrelevant clamped feedback experiment (E4) (Figure 3-1; analysis 5). However, a significant positive correlation only existed when the perturbation direction was congruent across the two experimental sessions. Finally, the rate of implicit adaptation in E4 was different between young and older adults (p = 0.002)(mean ± std; young: 8.2 × 10^−4^ ± 1.4 × 10^−4^, old: 16.9 × 10^−4^ ± 2.6 × 10^−4^) (analysis 5). In summary, this experiment provides evidence that implicit adaptation does not decrease with age and might even be increased in some conditions in older people. Further interpretation of these results is presented in the discussion section.

### 3.3. Explicit strategy is decreased with aging

Besides implicit recalibration, explicit strategy is another component that contributes to the overall adaptation (Mazzoni and Krakauer, 2006; Taylor et al., 2010; Morehead et al., 2015b). Here, explicit strategy was quantified with the cued motor adaptation experiment (Figure 1A; E1a and b) as the explicit adaptation level in each of the learning blocks.

In the first learning block, explicit adaptation level was significantly decreased for older compared to younger in experiment E1a, but not in experiment E1b (analysis 6) [Figure 4A-B; E1a: F(1,36) = 39.2, p < 10^−6^, 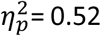; E1b: F(1,58) = 1.5, p = 0.2, 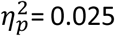]. In the second learning block, explicit adaptation level was significantly decreased for older compared to younger in E1a but not in E1b (analysis 6) [Figure 4A-B; E1a: F(1,36) = 64.6, p < 10^−8^, 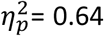; E1b: F(1,58) = 2.3, p = 0.1, 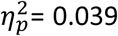]. For both learning blocks, no interaction between age and rotation direction was observed for E1a and E1b.

**Figure 4:**
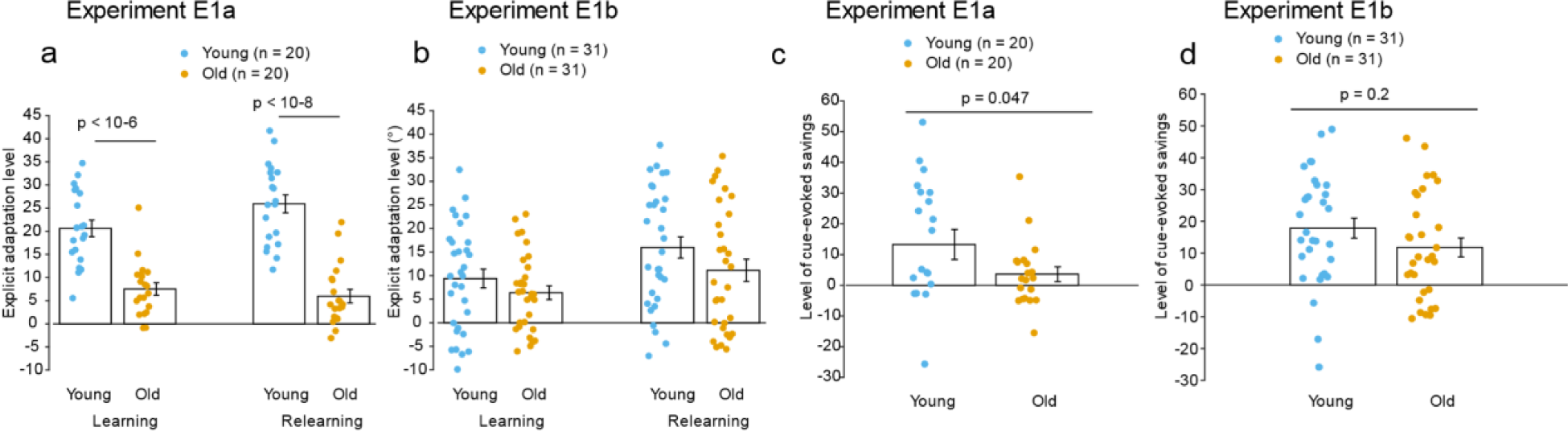
Decreased explicit strategy with aging. **A)** Significant decrease of explicit adaptation measured with E1a. **B)** Non-significant decrease of explicit adaptation with E1b. C) E1a: Significant decrease of cue-evoked savings for older compared to younger adults. **D)** E1b: Non-significant decrease of cue-evoked savings for older compared to younger adults.

In addition, we quantified savings as it has been linked to the retrieval of the explicit strategy (Morehead et al., 2015b). Therefore, we reasoned that savings should be smaller in old compared to young participants. Savings is quantified via the explicit adaptation level in the second learning block (Fig. 4A-B) and via cue-evoked savings (Figure 4C-D).

Savings can be quantified as the increase of explicit adaptation from the first towards the second learning block (analysis 6). An effect of learning block did not exist in E1a [E1a: F(1,38) = 1.9, p = 0.2] but the increase of explicit adaptation from first to second learning block was lower for older adults [E1a: interaction age*learning block: F(1,38) = 6.3, p = 0.02]. However, in E1b an effect of learning block existed [E1b: F(1,60) = 21.1, p < 10^−4^] but the increase of explicit adaptation from first to second learning block was not different between young and older adults [E1b: interaction age*learning block: F(1,60) = 0.5, p = 0.5].

Cue-evoked savings, which is the difference in adaptation level between the first trial of the relearning block and the first trial of the learning block, was significantly decreased for older compared to younger adults in E1a (analysis 7) [Figure 4C; E1a: F(1,35) = 4.25, p = 0.047, 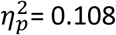], but not for E1b [Figure 4D; E1b: F(1,58) = 2.06, p = 0.2, 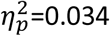].

All measures of explicit strategy were significantly decreased for the group of older participants in E1a, while in E1b this decrease was not significant. It might be tempting to explain this as questionable evidence for decline of explicit strategy with aging because E1b was the improved version of E1a and in E1b we didn’t observe a significant decline. However, after carefully observing the levels of explicit strategy (Figure 4A-B), it is clear that the main difference is the level of strategy-use of the younger participant which was lower in E1b compared to E1a. The level of explicit strategy of the older ones almost didn’t change. This different level of explicit strategy for younger participants in E1a and E1b can simply arise from natural sampling variability. Furthermore, also the overall adaptation level of younger adults (Figure 2A-B) was lower in E1b compared to E1a. As a consequence, overall adaptation was significantly decreased with aging in E1a, while in E1b overall adaptation was not significantly decreased. Altogether, these results show that explicit adaptation is decreased with aging, at least if overall adaptation is decreased as well. This suggests that differences in adaptation between young and old, if any, are due to the explicit component of adaptation. Therefore, we can predict that the amount of explicit component will be positively correlated with the amount of overall adaptation.

### 3.4. Overall adaptation is associated positively with explicit adaptation but negatively with implicit adaptation

Such positive correlation (Figure 5A-B; analysis 9) was indeed observed between overall adaptation and explicit adaptation while there was no or a negative correlation (Figure 5C-D) between overall and implicit adaptation. This suggests that the implicit component cannot be responsible for the decline of overall adaptation in elderly. The negative correlation between implicit and explicit adaptation (Figure 5E-F) may even suggest that the implicit component compensates for declines of explicit adaptation in elderly. We quantified the correlations by comparing components from different learning blocks in order to make sure that the measurements were independent from one another. Robust linear regression was used to verify that associations were resistant to age-effects (Fig. 5A-F), which seems indeed the case as beta values for adaptation components were significant for all associations except for one (Fig 5C).

**Figure 5:**
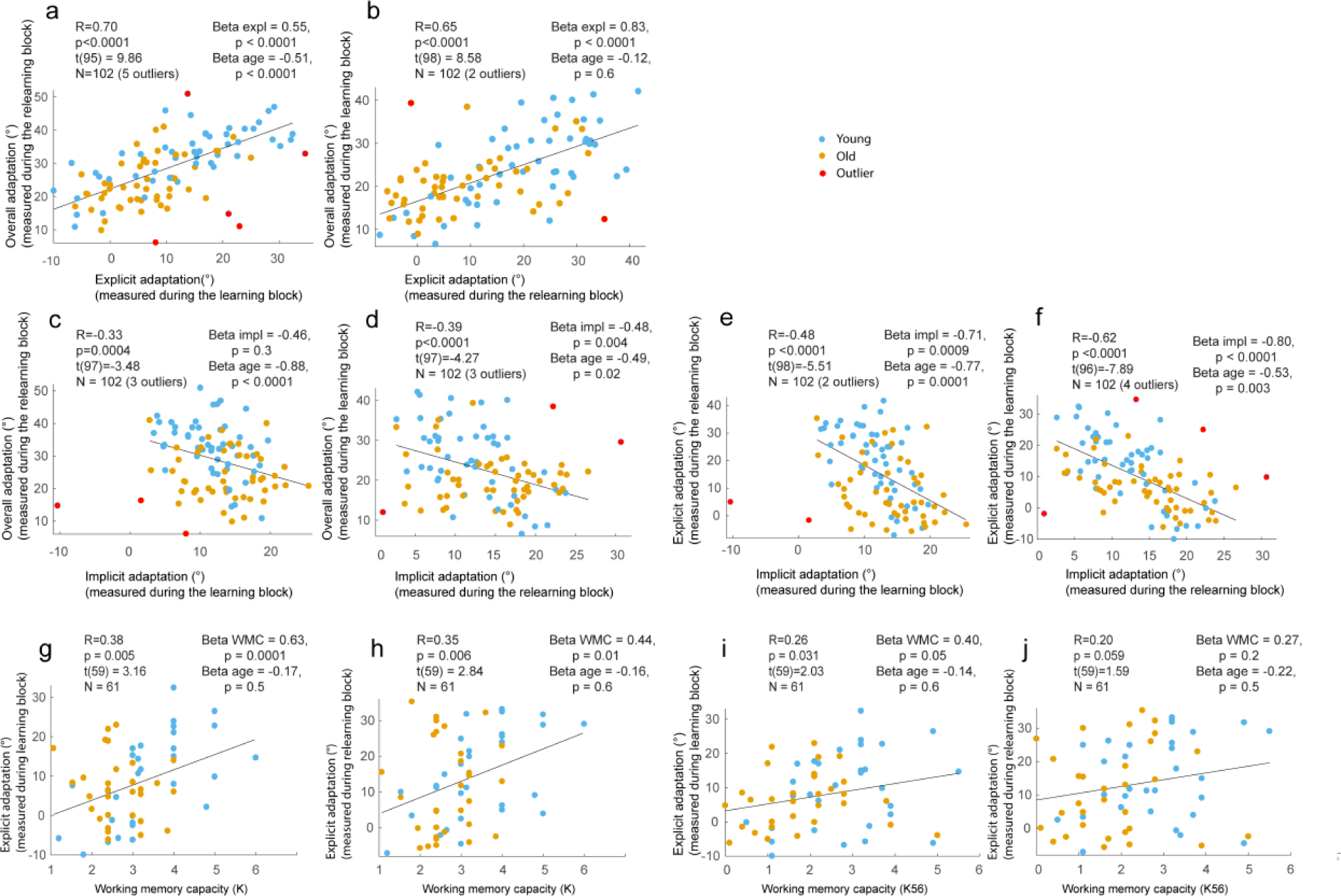
Explicit adaptation correlates positively with overall adaptation. Implicit adaptation correlates not or negatively with overall adaptation. Data were obtained with E1a and E1b with 102 young and elderly combined. **A-B)** Positive correlations between explicit adaptation and overall adaptation in learning and relearning blocks. **C-D)** No or negative correlations between implicit and overall adaptation in learning and relearning blocks. **E-F)** Negative correlations between implicit and explicit adaptation in learning and relearning blocks. **G-H)** Positive significant correlations between explicit strategy (y-axes) and visuospatial working memory capacity (K) (x-axes). **I-J)** Positive correlations (only 1 significant) between explicit strategy (y-axes) and visuospatial working memory capacity (K56) (x-axes). Pearson correlations were applied in A-F and Spearman correlations in G-J. All p-values of correlations are adjusted p-values with FDR. Beta values of linear regression are reported as well. See also Figure 5-1, 5-2 and 5-3.

In addition, all participants’ visuospatial working memory capacities (WMC) and neuropsychological status were quantified (analysis 10 and 11). From these measures it could be explored which broader cognitive domains are linked to the deficits in explicit strategy. WMC was declined in our older compared to our younger group (WMC-K: F(1,59) = 13.33, p = 0.0006; WMC-K56: F(1,59) = 9.95, p = 0.0025). In general, the younger group performed better than the older group for the RBANS cognitive assessment (F(1,60)= 37.6, p < 10^−7^). Explicit strategy correlated positively with visuospatial WMC when combining data from young and older participants (Figure 5G-J; analysis 12) but we found no correlation between WMC-K56 and explicit adaptation for young participants (Figure 5-1). RBANS task scores that might be related to explicit adaptation were selected and several scores correlated positively with explicit adaptation measures (Table 2; Figure 5-2; analysis 12) when combining data from young and old of the cued motor adaptation experiment (E1b). Correlations existed between explicit strategy and three RBANS cognitive scores (digit span, figure recall and line orientation test scores). Robust linear regression indicated that the correlations between explicit strategy and WMC-K and between explicit strategy and line orientation were not due to between-group differences. Together, this suggests that explicit strategy is positively associated with visuospatial abilities (visuospatial WMC and line orientation) in both younger and older adults. Finally, the balance between explicit and implicit adaptation was quantified (Figure 5-3, analysis 8) but was not altered with aging. However, this measurement was highly impacted by the variability of implicit adaptation in the denominator.

**Table 2:** Results of correlation analyses between adaptation components and working memory capacity or cognitive measures. Significant Spearman correlations existed between the explicit component (learning or relearning) and WMC-K and between the explicit component and three cognitive measures (i.e. digit span, figure recall and line orientation). Robust linear regression reveals that the correlations between explicit adaptation in relearning block and figure recall and between explicit adaptation and line orientation score are resistant to age-effects.

| Cognitive measure | Difference young-old (Mean ± StdErr) | Spearman correlation | | Robust linear regression (beta-coefficient and p-value for cognitive and age factors) | |
| --- | --- | --- | --- | --- | --- |
|  |  | Explicit learning | Explicit relearning | Explicit learning | Explicit relearning |
| WMC-K | 0.70 ± 0.23; p = 0.004 | R = 0.36; T(59) = 2.95, P = 0.01 | R = 0.35; T(59) = 2.87, P = 0.01 | Beta<sub>WMC</sub> = 0.63; P = 1e-04; Beta<sub>AGE</sub> = -0.17; p = 0.5 | Beta<sub>WMC</sub> = 0.44; P = 0.01; Beta<sub>AGE</sub> = -0.16; p = 0.6 |
| WMC-K56 | 0.98 ± 0.30; p = 0.002 | R = 0.26; T(59) = 2.09, P = 0.05 | R = 0.22; T(59) = 1.72, P = 0.09 | Beta<sub>WMC</sub> = 0.40; P = 0.05; Beta<sub>AGE</sub> = -0.14; p = 0.6 | Beta<sub>WMC</sub> = 0.27; P = 0.2; Beta<sub>AGE</sub> = -0.22; p = 0.5 |
| Figure copy | -0.03 ± 0.39, p = 0.9 | R=0.14; T(60)=1.05, p = 0.3; (p adj = 0.4) | R=0.24; T(60)=1.87, p =0.07; (p adj = 0.13) | Beta<sub>FIGC</sub> = 0.11; P = 0.6; Beta<sub>AGE</sub> = -0.33; p = 0.2 | Beta<sub>FIGC</sub> = 0.22; P = 0.3; Beta<sub>AGE</sub> = -0.40; p = 0.1 |
| Digit span | 1.39 ± 0.59, p = 0.02 | R=0.35; T(60)=2.87, p < 0.01; (p adj = 0.03) | R=0.27; T(60)=2.18, p =0.03; (p adj = 0.09) | Beta<sub>DIG</sub> = 0.03; P = 0.9; Beta<sub>AGE</sub> = -0.20; p = 0.5 | Beta<sub>DIG</sub> = 0.07; P = 0.7; Beta<sub>AGE</sub> = -0.27; p = 0.3 |
| Coding total | 10.94 ± 1.95, p < 1x10<sup>-06</sup> | R=0.13; T(59)=1.04, p =0.3; (p adj = 0.4) | R=0.17; T(60)=1.29, p =0.2; (p adj = 0.3) | Beta<sub>COD</sub> = 0.37; P = 0.08; Beta<sub>AGE</sub> = -0.23; p =0.5 | Beta<sub>COD</sub> = 0.41; P = 0.06; Beta<sub>AGE</sub> = -0.38; p = 0.3 |
| Figure recall | -0.03 ± 0.39, p = 0.01 | R=0.15; T(59)=1.13, p =0.3; (p adj = 0.4) | R=0.38; T(60)=3.17, p < 0.01; (p adj = 0.02) | Beta<sub>FIGR</sub> = 0.07; P = 0.8; Beta<sub>AGE</sub> = -0.30; p = 0.3 | Beta<sub>FIGR</sub> = 0.32; P = 0.2; Beta<sub>AGE</sub> = -0.21; p = 0.5 |
| Line orientation | 0.23 ± 0.42, p = 0.6 | R=0.33; T(55)=2.67, p < 0.01; (p adj = 0.03) | R=0.60; T(56)=5.70, p < 10<sup>-6</sup>; (p adj < 0.0001) | Beta<sub>LO</sub> = 0.34; P =0.06; Beta<sub>AGE</sub> =-0.32; p = 0.2 | Beta<sub>LO</sub> = 0.48; P = 0.003; Beta<sub>AGE</sub> = -0.31; p = 0.2 |

### 3.5. Short-term relative retention of motor adaptation memory is not affected

In the retention hypothesis, a decreased trial-to-trial retention of motor adaptation is thought to account for a reduced amount of adaptation with aging (Trewartha et al., 2014; Malone and Bastian, 2015). To investigate the age-effect on retention of motor memory, we introduced one-minute breaks as these short breaks are known to capture the dynamics of motor adaptation well because most of the decay of motor adaptation occurs in this short timescale (Hadjiosif and Smith, 2013). Short-term relative retention was assessed with a regular visuomotor rotation experiment with three breaks of one minute in each learning block (E3; Figure 1C) and with three breaks of one minute in the task-irrelevant clamped feedback experiment (E4; Figure 1D). Results are shown in Figure 6.

**Figure 6:**
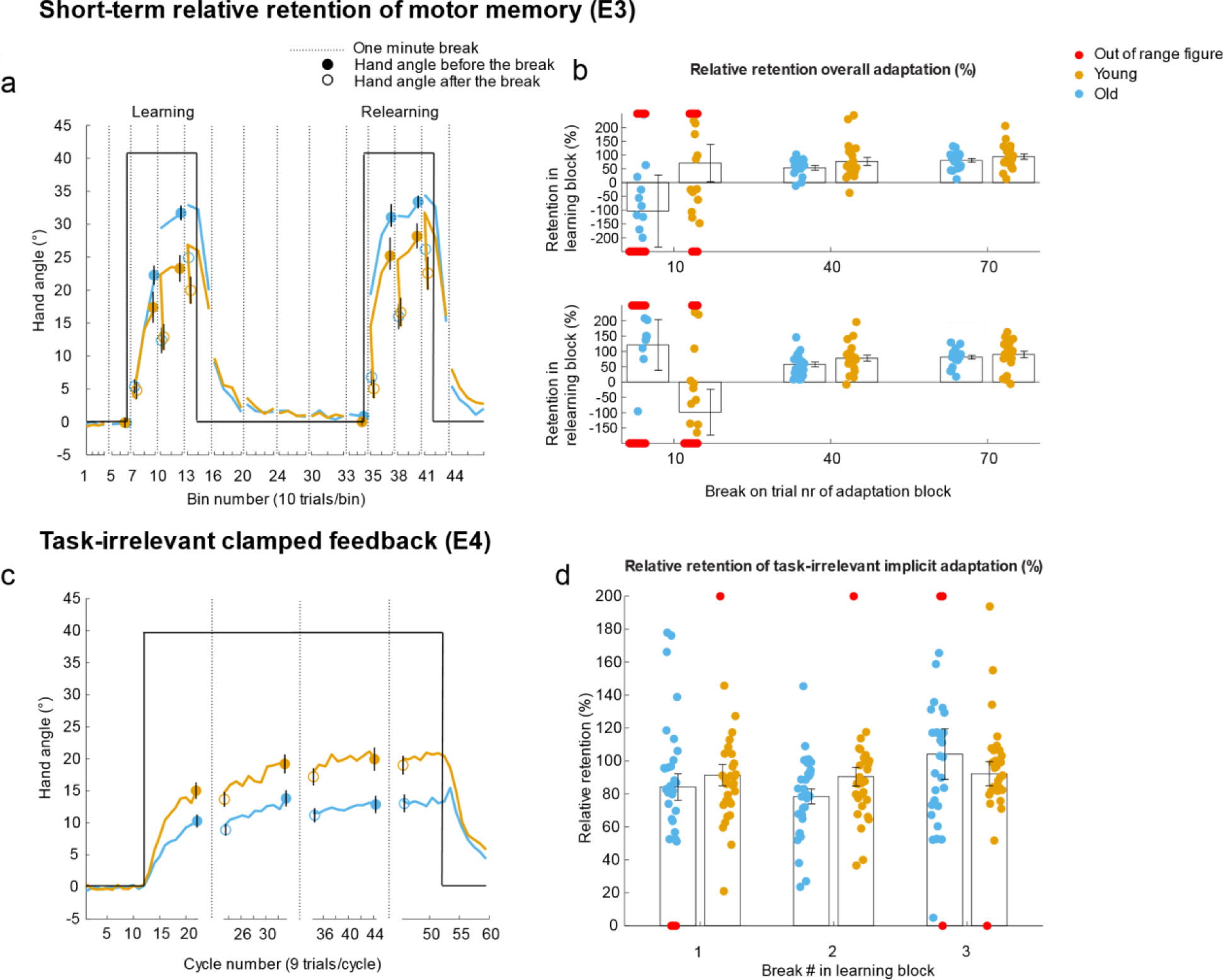
Relative retention of motor adaptation. **A)** Decreased overall adaptation in older vs younger adults in E3. Blue and orange lines represent the average learning curves for young and older adults respectively. The blue and orange dots represent the overall adaptation level before the three breaks in each adaptation block. The blue and orange unfilled dots represent the remaining adaptation level after a one-minute break, i.e. stable component of adaptation. **B)** Relative retention of motor adaptation memory was not different for young and old in E3 for each of the three breaks in both learning blocks. **C)** Increased adaptation for older vs younger adults during task-irrelevant clamped feedback (E4). **D)** Relative retention of motor adaptation memory was not different for young and old in E4 for each of the three breaks during the task-irrelevant clamped feedback.

Short-term relative retention was not significantly different for older compared to younger adults (analysis 13) [E3: F(1,35) = 2.2, p = 0.14, 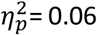; E4: F(1,58) = 0.1, p = 0.7, 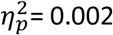]. The visuomotor rotation experiment (E3) consisted of two learning blocks. Relative retention was not different for the two learning blocks [F(1,35) = 0.26, p = 0.6]. Each learning block consisted of three one-minute breaks. The relative retention was not significantly different for the different breaks in experiment E3 and E4 [E3: F(2,70) = 2.7, p = 0.07; E4: F(2,116) = 1.3, p = 0.3]. Given that the data for the first breaks in E3 are extremely noisy, we redid the analyses on the last two breaks only. These analyses confirmed that there were no differences in relative retention between the two age groups [F(1,35) = 2.6, p = 0.11, 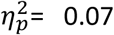]. Finally, no interactions between age and other factors were observed. Therefore, we can conclude that short-term relative retention was not affected with aging.

### 3.6. Reaction time not a good indicator for explicit strategy in older adults

In young adults higher reaction times are associated with more explicit strategy (Fernandez-ruiz et al., 2011; Taylor and Mcdougle, 2016). However, across groups this relation was not present in our data because we found a relative increase in reaction times (RT) despite a reduced explicit strategy for older adults instead of a decrease of RT for older adults compared to younger ones (Figure 7) (analysis 14) (E1a: learning: p = 0.1; relearning: p = 0.02; E1b: learning: p = 0.05; relearning: p = 0.3; E3: learning: p = 0.02; relearning: p = 0.01; E4: p = 0.02).

**Figure 7:**
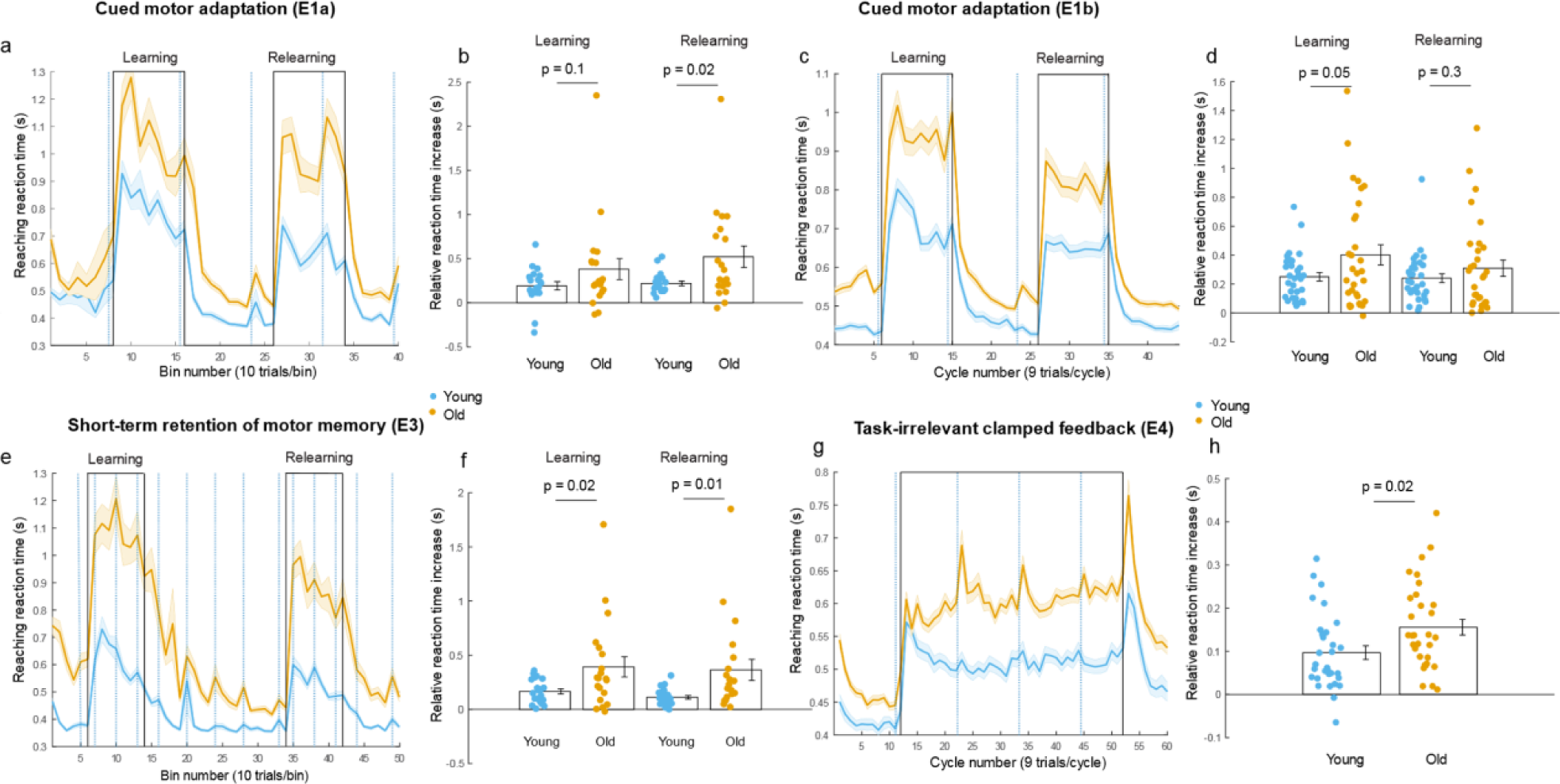
Reaction time (RT) during the different adaptation experiments. During learning blocks RT always increased. However, older adults increased their RT more than younger adults during adaptation. **A)** RT during cued adaptation experiment (E1a), **B)** Relative RT increase during learning blocks of the cued motor adaptation experiment (E1a), **C)** RT during cued adaptation experiment (E1b). **D)** Relative RT increase during learning blocks of the cued motor adaptation experiment (E1b), **E)** RT during retention experiment (E3), **F)** Relative RT increase during learning blocks of the short-term retention experiment (E3), **G)** RT during task-irrelevant clamped feedback adaptation (E4). **H)** Relative RT increase during task-irrelevant clamped feedback adaptation (E4).

In addition to the effect of age on reaction time, other experimental variables were investigated with respect to age such as reaching time, movement curvature, etc. In general, older adults were reaching slower towards the targets and they took more time between the different reaching trials than young adults. In addition, their reaching movements were more curved and they obtained lower scores. For completeness, we presented the different variables in Table 3. We do not think that these variables influenced our adaptation results because they could not account for the fact that some components of adaptation are improved or unimpaired in older participants (e.g. Implicit component and relative retention) while others are affected (e.g. explicit component, overall adaptation). In addition, for every reaching trial the adaptation level was measured before any visual feedback adjustments could be applied by the participant which implies that increased reaching times for elderly will not impact our measures of adaptation components.

**Table 3:** Secondary analyses of reaching behavior with nine variables, comparison between young and older adults.

|  | E1a |  | E1b |  | E2 |  |
| --- | --- | --- | --- | --- | --- | --- |
| | Mean difference ± SEM<br>(young-old) | Statistics | Mean difference ± SEM<br>(young-old) | Statistics | Mean difference ± SEM<br>(young-old) | Statistics |
| Trial time duration (s) | -0.66 ± 0.20 | P = 0.002;<br>T(38) = -3.33 | -0.37 ± 0.11 | P = 0.00098;<br>t(60) = -3.47 | -0.195 ± 0.068 | P = 0.0055;<br>t(79) = -2.85 |
| Reaching time (s) | -0.046 ± 0.018 | P = 0.016;<br>T(38) = -2.52 | -0.010 ± 0.003 | P = 0.00046;<br>t(60) = -3.71 | -0.038 ± 0.009 | P = 1.26e-04;<br>t(79) = -4.03 |
| Max displacement (mm) | -0.8 ± 0.2 | P = 1.54e-04;<br>T(38) = -4.20 | -0.6 ± 0.2 | P = 0.003;<br>t(60) = -3.13 | -0.36 ± 0.011 | P = 0.0014;<br>t(79) = -3.30 |
| Surface displacement (dm<sup>2</sup>) | -0.0034 ± 0.0010 | P = 0.002;<br>T(38) = -3.38 | -0.005 ± 0.001 | P = 0.0002;<br>t(60) = -3.92 | -0.0020 ± 0.0007 | P = 6.45e-05;<br>t(60) = -4.29 |
| Number of too fast trials | -9.15 ± 4.33 | P = 0.041;<br>T(38) = -2.11 | 1.387 ± 4.041 | P = 0.008;<br>t(60) = -2.76 | 0.274 ± 0.690 | P = 0.69;<br>t(79) = 0.40 |
| Number of too slow trials | -23.0 ± 6.8 | P = 0.002;<br>T(38) = -3.40 | -27.5 ± 5.8 | P = 1.23e-05;<br>t(60) = -4.77 | -25.7 ± 5.2 | P = 4.85e-06;<br>t(79) = -4.91 |
| Total score | 5941.3 ± 815.0 | P = 1.00e-08;<br>T(38) = 7.29 | 3279.7 ± 903.1 | P = 0.00058;<br>t(60) = 3.63 | 2033.6 ± 429.3 | P = 9.39e-06;<br>t(79) = 4.74 |
| Inter-trial interval (s) | -0.691 ± 0.204 | P = 0.002;<br>T(38) = -3.38 | -0.378 ± 0.116 | P = 0.0019;<br>t(60) = -3.25 | -0.172 ± 0.078 | P = 0.03;<br>t(79) = -2.20 |
| Experiment duration (s) | -372.75 ± 103.35 | P = 8.89e-04;<br>T(38) = -3.61 | -290 ± 69 | P = 9.74e-05;<br>t(60) = -4.18 | -106.85 ± 40.96 | P = 0.01;<br>t(79) = -2.61 |
|  | E3 |  | E4 |  |  |  |
| | Mean difference ± SEM<br>(young-old) | Statistics | Mean difference ± SEM<br>(young-old) | Statistics | | |
| Trial time duration (s) | -0.717 ± 0.204 | P = 0.001; t(39) = -3.51 | -0.39 ± 0.09 | P = 7.99e-05; t(60) = -4.23 | | |
| Reaching time (s) | -0.073 ± 0.019 | P = 5.55e-4; t(39) = -3.76 | -0.015 ± 0.004 | P = 4.63e-04; t(60) = -3.70 | | |
| Max displacement (mm) | -0.006 ± 0.002 | P = 0.003; t(39) = -3.16 | -0.006 ± 0.001 | P = 7.05e-07; t(60) = -5.54 | | |
| Surface displacement (dm<sup>2</sup>) | -0.004 ± 0.001 | P = 0.01; t(39) = -2.77 | -0.0030 ± 0.0006 | P = 1.67e-05; t(60) = -5.06 | | |
| Number of too fast trials | -13.69 ± 5.51 | P = 0.017; t(39) = -2.49 | -0.226 ± 4.331 | P = 0.96; t(60) = -0.05 | | |
| Number of too slow trials | -22.7 ± 7.7 | P = 0.005; t(39) = -2.95 | -16.32 ± 5.44 | P = 0.004; t(60) = -3.00 | | |
| Total score | 8494.4 ± 2205.5 | P = 4.26e-04; t(39) = 3.85 | 2161.6 ± 697.99 | P = 0.003; t(60) = 3.10 | | |
| Inter-trial interval (s) | -0.778 ± 0.227 | P = 0.002; t(39) = -3.42 | -0.393 ± 0.095 | P = 0.0001; t(60) = -4.12 | | |
| Experiment duration (s) | -7.97 ± 2.25 | P = 0.001; t(39) = -3.54 | -229 ± 53 | P = 5.23e-05; t(60) = -4.36 | | |

## 4. Discussion

In this paper, we tested three different hypotheses that could account for the observed age-related deficits in motor adaptation tasks in several experiments over large groups of participants. First, we found that implicit adaptation was intact or even increased in older adults compared to younger ones. This is inconsistent with the internal model hypothesis. Second, our data are consistent with the strategy hypothesis as we found a lower explicit component during motor adaptation in older people. Third, we did not find any support for the retention hypothesis, as short-term retention of motor memory was indistinguishable between young and old participants. Together, these results suggest that, despite age-related cerebellar degeneration, internal model function remains intact with aging and could even compensate for deficits in the explicit component.

### 4.1. Internal model recalibration is intact or even increased in older people

While previous studies assumed that age-related degeneration of the cerebellum was responsible for the decline of motor adaptation with aging (Seidler, 2006, 2007), our data are at odds with this view. Indeed, across four experiments (Figure 3), we found that internal model recalibration was intact or even increased with aging. Rather, this is consistent with the fact that several studies did not observe differences in after-effect between young and old people (Fernández-Ruiz et al., 2000; Buch et al., 2003; Heuer and Hegele, 2008, 2014; Hegele and Heuer, 2010; Sombric et al., 2017). Yet, our data support the possibility that internal model function could be increased in older people as we found that both error-sensitivity and implicit adaptation increased with aging. If this function is increased, it could demonstrate the interplay between different brain regions, where one brain area (here the cerebellum) might compensate for deficits in another brain region. This is consistent with brain imaging studies where the cerebellum is increasingly activated by motor tasks in old but not in young people (Mattay et al., 2002; Heuninckx et al., 2005, 2008) and with the fact that cerebellar volume is shrinking relatively less with age compared to other brain regions such as lateral prefrontal cortex (Raz et al., 2005). In addition, we observed intact internal model function despite the known age-related changes in cerebellar structure. While we don’t currently have data on our participants about the extent of cerebellar degeneration, decrease in cerebellar volumes between 30 and 70 years old appears to be consistent across individuals (Walhovd et al., 2011). If so, our study suggests an important dissociation between structure and function of the anterior part of the cerebellum responsible for motor control (Schmahmann, 2018) and raises the question of how much volume of the anterior lobe can be lost without affecting internal model recalibration. Indeed, it is known that extensive loss of cerebellar volumes in patients with cerebellar degeneration affects internal model function (Smith and Shadmehr, 2005; Morton and Bastian, 2006; Taylor et al., 2010; Gibo et al., 2013; Morehead et al., 2017; Criscimagna-hemminger et al., 2018). In our sample, age-related changes of the cerebellum, if any, had no impact on internal model recalibration during visuomotor adaptation. Below, we outlined three possible explanations for these observations.

A first explanation for the observation of increased error-sensitivity and implicit adaptation is an altered sensory integration with aging and an intact internal model recalibration. If visual acuity is similar for both young and older participants and if arm proprioception is reduced in older participants (Goble et al., 2009), then the weighting of visual compared to proprioceptive feedback might be increased in older participants following Bayesian sensorimotor integration(Ernst and Banks, 2002; Van Beers et al., 2002; Kording and Wolpert, 2004). The up weighted visual feedback would create an increased sensory-prediction error for older versus younger adults and as such result in an increased reaction to error in the single trial error learning experiment (E2) and a higher level of asymptote in the task-irrelevant feedback experiment (E4). Hence, this first explanation is consistent with a change in the input to the internal model with aging and requires intact internal model function.

The second explanation is an increased weighting of the predicted sensory feedback from the internal model with respect to the sensory feedback because of the age-related reduction in proprioceptive acuity. Indeed, it is known that state estimation results from a weighted average between sensory information and “prior” predictions in function of their reliability (Körding and Wolpert, 2004). Changes in reliability of one of those two signals could therefore impact motor control by altering the weighting of these two signals. Recently, such an increased reliance on sensorimotor prediction in proportion to reduced proprioceptive sensitivity was observed with aging for a force matching task (Wolpe et al., 2016). Therefore, we proposed here that internal models are recalibrated similarly in young and elderly people. However, because of the age-related decrease in the reliability of the proprioceptive information, the output of the internal model has more weight in the reliability weighted integration between internal model output and proprioceptive information.

A third explanation that could account for the observed increased error-sensitivity and implicit adaptation in elderly people is an actual increase of internal model recalibration with aging. An increased internal model recalibration means that older adults react more to sensory prediction errors of the same amplitude compared to younger people. This explanation would be compatible with the push-pull mechanism between explicit and implicit adaptation proposed by several authors (Taylor and Ivry, 2011; Heuer and Hegele, 2014; Christou et al., 2016). In these studies, the authors suggest that the decrease in explicit adaptation elicit the increase in implicit adaptation (i.e. internal model recalibration) in order to compensate for it. This is also compatible with our data as we observed a negative correlation between explicit and implicit adaptation components (Figure 5I-J). However, this explanation can only hold when both an explicit and implicit component of motor adaptation are present like in the cued motor adaptation experiment (E1). Yet, we also observed increased implicit adaptation in the absence of this impaired explicit component in the single-trial learning task (E1) and in the task-irrelevant clamped feedback task (E4). Therefore, our data suggest that increased internal model recalibration is independent of the deficit in the explicit component of adaptation, at least in some situations. In sum, the third explanation suggests that the same input results in an increased output by increased internal model recalibration.

These different explanations are compatible with each other, as such they might all be valid. In addition, they all rely on the assumption that internal model function is not deteriorated by aging. Future experiments should be designed to test each of them separately. By doing so, we can eventually build a model for sensorimotor integration that is valid across age.

Finally, it is worth noting that our measure of implicit adaptation could be slightly overestimated in the cued motor adaptation. Indeed, it has been shown in such experiments that the adaptation generalizes around the aiming direction (Day et al., 2016; McDougle et al., 2017). Given that older adults changed their aiming direction less, generalization of implicit adaptation occurs closer to the probed target and might yield a spurious increase in our estimation of the implicit component. However, it cannot explain the increased error-sensitivity (E2) and implicit adaptation (E4).

### 4.2. The cognitive component of adaptation is reduced in elderly people

We observed a difference in overall motor adaptation with aging when there was an age-related deficit in explicit adaptation (Figure 4A-B). In contrast, when the explicit component was unimpaired, so was the overall motor adaptation, as observed in experiment E1b. This observation provides additional evidence for the strategy hypothesis which states that explicit adaptation is affected with aging and is causing the decline of overall motor adaptation (Bock, 2005; Heuer and Hegele, 2008; Hegele and Heuer, 2010). In addition to a decline in explicit adaptation, we found a reduction of savings with aging. First, the increase of explicit adaptation level from the first to the second learning block was lower in older adults. This finding is in contrast with previous studies that reported no difference in savings with aging (Seidler, 2007). Second, cue-evoked savings (Morehead et al., 2015b), during which participants adapted within a single trial on the basis of a contextual cue, was lower in older adults (Figure 4C and D). Across these measures of adaptation, we found a deficit in explicit component with age.

It is currently unknown which brain area is critical for the explicit component of motor adaptation. Yet, it has been suggested that the cerebellum might play a role in the formation of an explicit strategy (Butcher et al., 2018) independently of it role in internal model recalibration. Indeed, cerebellar patients with posterior lobe lesions develop cognitive deficits (cerebellar cognitive affective syndrome), while anterior lobe lesions result in motor deficits (Schmahmann, 2018). Therefore, cerebellar degeneration might still be involved in decline of motor adaptation with aging because it is linked to deficits in explicit strategy and not because the cerebellum is crucial for internal model recalibration (internal model hypothesis). Future neuro-imaging studies should investigate the link between anterior and posterior cerebellar volume and age-related deficits in motor adaptation.

### 4.3. Short-term retention of motor memories is intact in elderly people

We observed no age-related decrease of relative retention of motor adaptation. This implies that the rate of retention is not affected with aging and that the retention hypothesis is not correct. Our observation of no short-term retention deficit after a one minute break is in contrast with previous work that shows retention deficits of motor adaptation after breaks of five minutes (Malone and Bastian, 2015) or that shows retention deficits with an indirect modeling approach (Trewartha et al., 2014). An explanation for this contradiction is the difference between the different paradigms (length of breaks, walking adaptation vs reaching adaptation, the indirect versus direct approach). Further research is required to explain these differences.

## 5. Conclusion and outlook

Our results show that neither internal model recalibration nor short-term retention of motor adaptation are reduced with aging. Rather, we demonstrate that cognitive processes involved in motor adaptation are impaired in older participants. Therefore, our data provides support for the “strategy hypothesis”, which states that age-related decline in motor adaptation is due to cognitive deficits. This study is the largest study so far investigating the impact of aging on motor adaptation processes. Yet, it leaves us with two important puzzles for the future. First, we expect a remarkable dissociation between age-related degeneration of the cerebellum and the intact or even improved function of this brain region (i.e. internal model function). Future studies should investigate the link between cerebellar volume and implicit motor adaptation. In addition, this study focuses on an older population between 60 and 75 years old, which is still relatively young. It remains to be investigated whether further reduction in cerebellar volume through aging elicit an impairment in internal model function for people above 80 years old. Second, possible explanation of the increased internal model recalibration observed are linked to proprioceptive acuity. Therefore, future studies should systematically investigate the link between implicit motor adaptation and error-sensitivity and proprioceptive acuity (Ostry et al., 2010; Wong et al., 2012). Finally, this study only covers some aspects of motor adaptation. It remains to be seen whether other processes underlying motor adaptation such as reinforcement learning (Huang et al., 2011; Izawa and Shadmehr, 2011; Orban de Xivry and Lefèvre, 2015) are affected by aging and whether the quality of motor memories, as probed by motor memory interference studies (Shadmehr and Holcomb, 1997; Krakauer et al., 2005; Nozaki et al., 2006, 2016), is similar in young and old participants.

Our study suggests that local structural brain changes might not always directly affect function and that neuroscience requires intelligent analyses and behavioral approaches in order to better understand how the brain works (Jonas and Kording, 2017; Krakauer et al., 2017; Herzfeld and Shadmehr, 2018). By following such an approach, we can eventually build models for sensorimotor functioning that are valid across age. Such models could help us to design smarter rehabilitation strategies for older adults with neurological disease or to delay deterioration of motor functioning with aging.

## Acknowledgments

This work was supported by an internal grant of the KU Leuven (STG/14/054) and by the FWO (1519916N). We thank Bradley R. King and Matthieu P. Boisgontier for discussing the results together and Robert Hardwick for his helpful comments on an earlier version of this manuscript.

## Extended Data

### Correlation between cued implicit adaptation (E1b) and task-irrelevant clamped feedback adaptation (E4)

**Figure 3-1:**
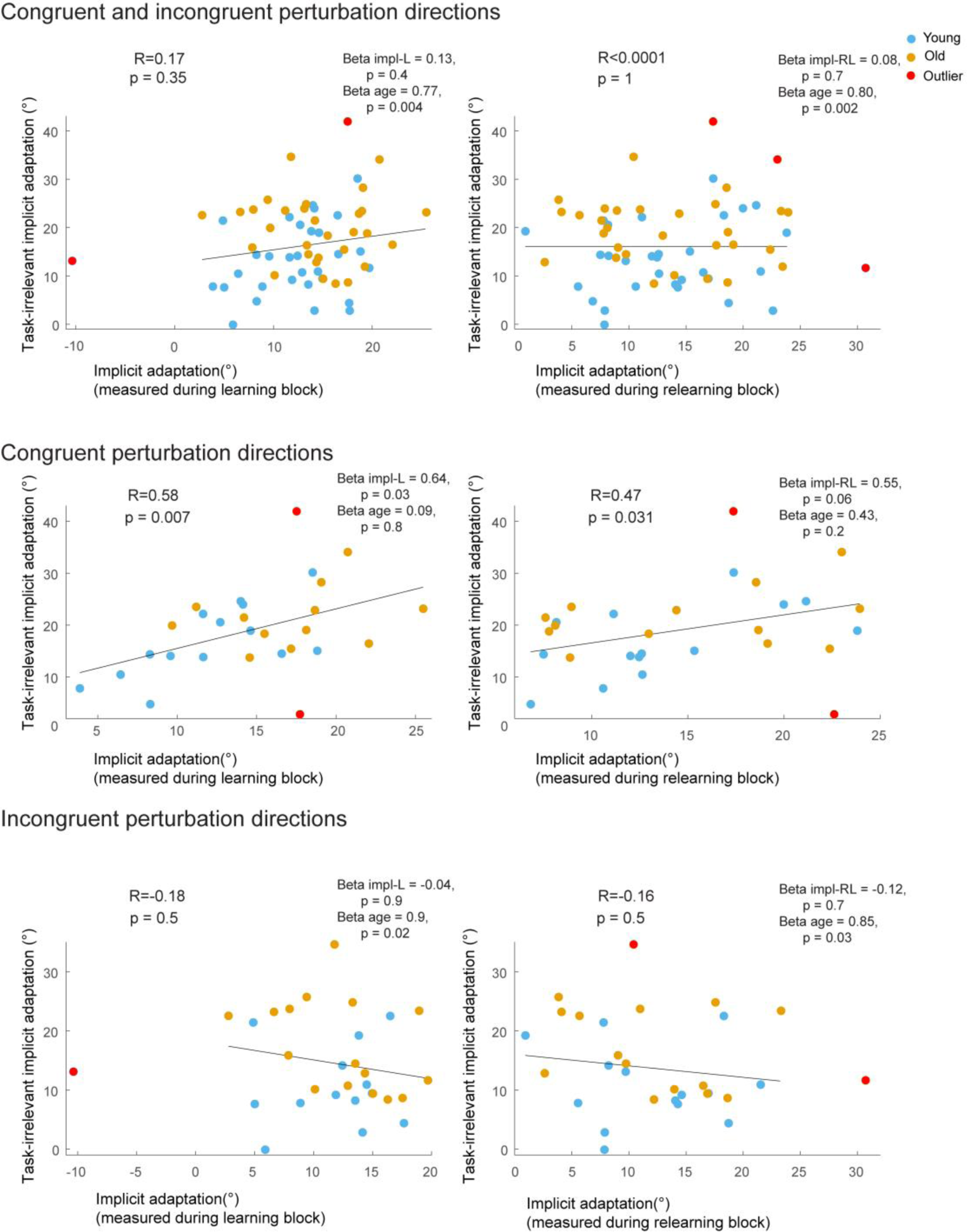
Correlation between cued implicit adaptation (E1b) and task-irrelevant clamped feedback adaptation (E4) (preregistered secondary analysis) (analysis 5). P-values adjusted with FDR. Beta values are obtained with robust linear regression.

### Failed replication of correlation between WMC-K56 and explicit adaptation

**Figure 5-1:**
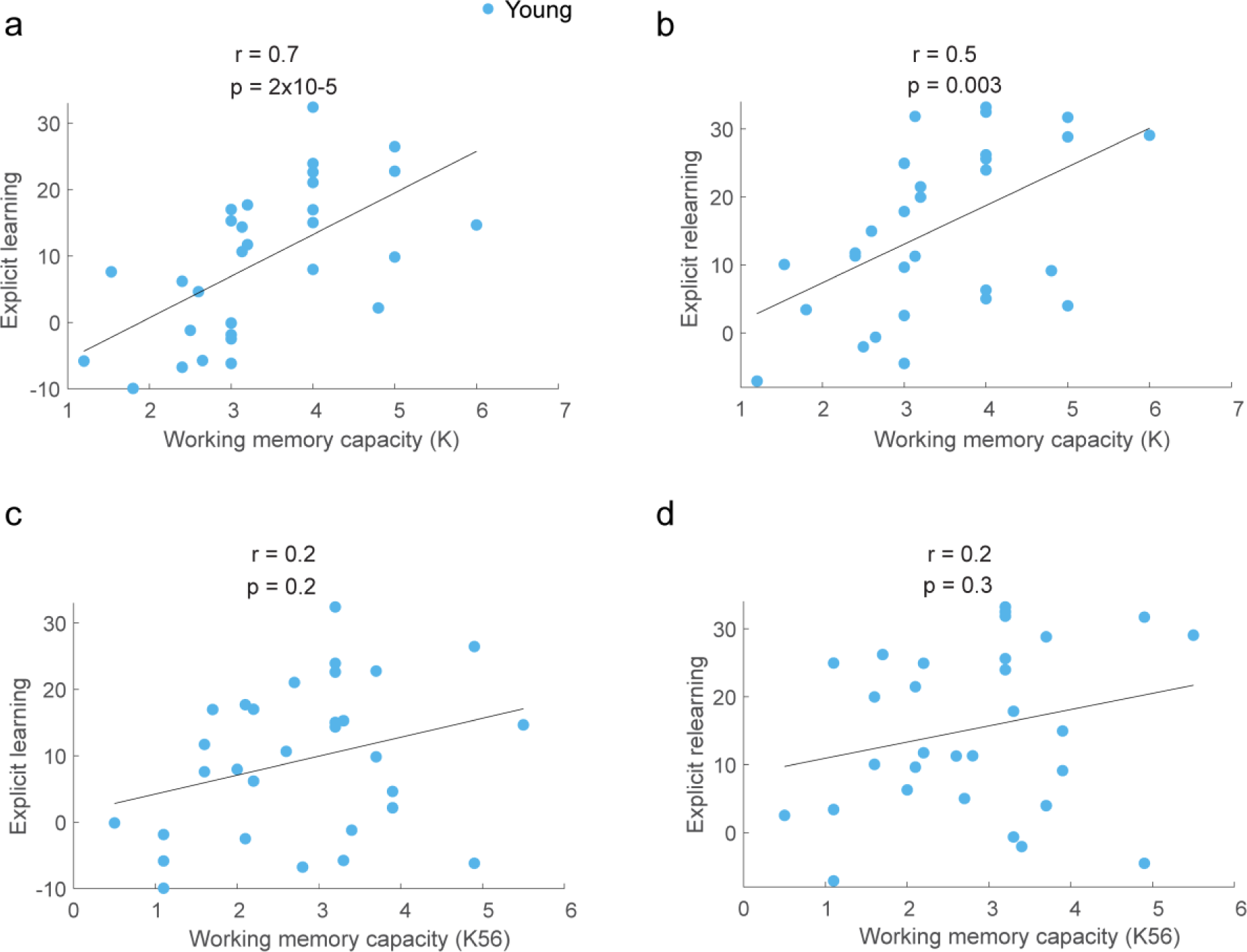
Failed replication of correlation between WMC-K56 and explicit adaptation. **A-B)** Significant correlation between working memory capacity (WMC-K) and explicit adaptation for young participants. **C-D)** Non-significant correlation between working memory capacity (WMC-K56) and explicit adaptation for young participants. WMC-K56 was calculated with five and six items, while WMC-K was calculated with all three to six items. Correlation coefficients were Spearman coefficients.

### Explicit strategy positively associated with RBANS scores

**Figure 5-2:**
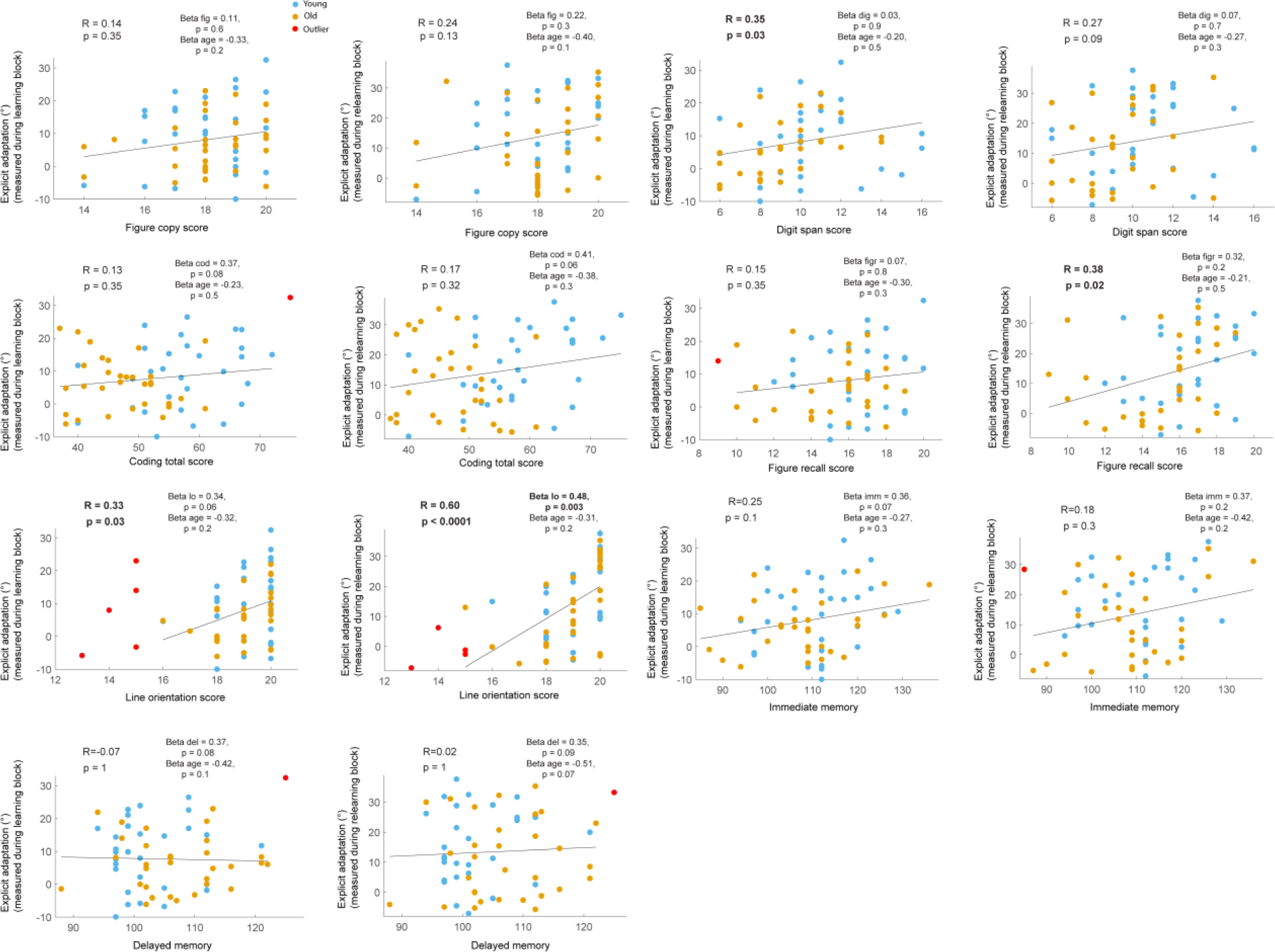
Explicit strategy positively associated with RBANS scores. Spearman correlations between raw RBANS test scores and explicit components of motor adaptation. Significant correlations (adjusted p-values with FDR) existed between the explicit component and three cognitive measures (i.e. digit span, figure recall and line orientation). Beta values of robust linear regression are reported as well, beta value for line orientation score remained significant.

### Balance between explicit and implicit adaptation unaltered with aging

**Figure 5-3:**
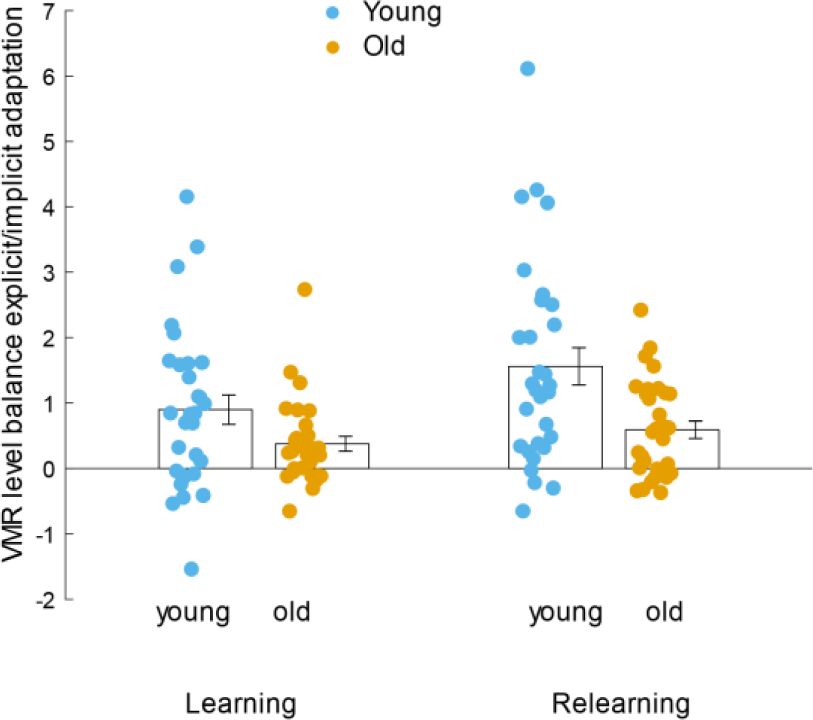
Balance between explicit and implicit adaptation unaltered with aging. Balance of explicit over implicit adaptation is not different in older compared to younger adults with the explicit component from the second cued motor adaptation experiment (E1b) and the implicit component from the task irrelevant-feedback experiment (E4) [F(1,53) = 1.6, p = 0.2, 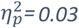 after removal of one outlier] (analysis 8). Congruency of rotation and rotation direction did not affect the balance [congruency effect: F(1,53)= 0.83, p = 0.4; rotation effect: F(1,53) = 1.4, p = 0.2]. The balance was increased in relearning compared to learning [F(1,53) = 5.3, p = 0.02]. No significant interactions between age and the other factors were observed. Contrary to expectations, no significant balance difference existed between young and older adults. The balance variable consisted of a division by the implicit adaptation which induced high variability. Due to the high variability the power of the experiment was probably too low to detect a significant difference.

## Notes

**Conflict of interest** The authors declare no competing financial interests.

## References

Anguera J a, Reuter-Lorenz P a, Willingham DT, Seidler RD (2011) Failure to engage spatial working memory contributes to age-related declines in visuomotor learning. J Cogn Neurosci 23:11–25.

Benjamini Y, Hochberg Y (1995) Controlling the False Discovery Rate: A Practical and Powerful Approach to Multiple Testing. 57:289–300.

Bernard JA, Seidler RD (2014) Moving forward: Age effects on the cerebellum underlie cognitive and motor declines. Neurosci Biobehav Rev 42:193–207.

Bock O (2005) Components of sensorimotor adaptation in young and elderly subjects. Exp Brain Res 160:259–263.

Bock O, Girgenrath M (2006) Relationship between sensorimotor adaptation and cognitive functions in younger and older subjects. Exp Brain Res 169:400–406.

Bock O, Schneider S (2002) Sensorimotor adaptation in young and elderly humans. Neurosci Biobehav Rev 26:761–767.

Boisgontier MP (2015) Motor aging results from cerebellar neuron death. Trends Neurosci 38:127–128.

Boisgontier MP, Nougier V (2013) Ageing of internal models: From a continuous to an intermittent proprioceptive control of movement. Age (Omaha) 35:1339–1355.

Buch ER, Young S, Contreras-Vidal JL (2003) Visuomotor Adaptation in Normal Aging. Learn Mem 10:55–63.

Butcher XPA, Ivry RB, Kuo S, Rydz D, Krakauer JW, Taylor XJA (2018) Higher Neural Functions and Behavior The cerebellum does more than sensory prediction error-based learning in sensorimotor adaptation tasks. :1622–1636.

Christou AI, Miall RC, McNab F, Galea JM (2016) Individual differences in explicit and implicit visuomotor learning and working memory capacity. Sci Rep 6:36633.

Criscimagna-hemminger SE, Bastian AJ, Shadmehr R (2018) Size of Error Affects Cerebellar Contributions to Motor Learning. :2275–2284.

Day KA, Roemmich RT, Taylor JA, Bastian AJ (2016) Visuomotor learning generalizes around the intended movement. eNeuro 3:1–12.

Duff K, Patton D, Schoenberg MR, Mold J, Scott JG, Adams RL (2003) Age-and Education-Corrected Independent Normative Data for the RBANS in a Community Dwelling Elderly Sample Age-and Education-Corrected Independent Normative Data for the RBANS in a Community Dwelling Elderly Sample. 4046.

Ernst M, Banks MS (2002) Humans integrate visual and haptic information in a statistically optimal fashion. Nature 415:429–433.

Fernández-Ruiz J, Hall C, Vergara P, Díaz R (2000) Prism adaptation in normal aging: Slower adaptation rate and larger aftereffect. Cogn Brain Res 9:223–226.

Fernandez-ruiz J, Wong W, Armstrong IT, Flanagan JR (2011) Relation between reaction time and reach errors during visuomotor adaptation. Behav Brain Res 219:8–14.

Fine MS, Thoroughman KA (2006) Motor Adaptation to Single Force Pulses?: Sensitive to Direction but Insensitive to Within-Movement Pulse Placement and Magnitude. :710–720.

Folstein M, Folstein S, McHugh P (1975) A practical state method for. 12:189–198.

Gibo TL, Criscimagna-Hemminger SE, Okamura AM, Bastian AJ (2013) Cerebellar motor learning: are environment dynamics more important than error size? J Neurophysiol 110:322–333.

Goble DJ., Coxon JP., Wenderoth N, Van Impe A, Swinnen SP (2009) Proprioceptive sensibility in the elderly: Degeneration, functional consequences and plastic-adaptive processes. Neurosci Biobehav Rev 33:271–278.

Hadjiosif AM, Smith M a. (2013) Savings is restriced to the temporally labile component of motor adaptation. Tcmc:1–2.

Haith AM, Huberdeau DM, Krakauer JW (2015) The Influence of Movement Preparation Time on the Expression of Visuomotor Learning and Savings. J Neurosci 35:5109–5117.

Hegele M, Heuer H (2010) The impact of augmented information on visuo-motor adaptation in younger and older adults. PLoS One 5:e12071.

Herzfeld DJ, Shadmehr R (2018) Impact of Human Behavioral Papers at Journal of Neuroscience. :1–7.

Hesterberg T, Monaghan S, Moore DS, Clipson A, Epstein R (2003) BOOTSTRAP METHODS AND PERMUTATION TESTS COMPANION CHAPTER 18 TO University of Puget Sound.

Heuer H, Hegele M (2008) Adaptation to visuomotor rotations in younger and older adults. Psychol Aging 23:190–202.

Heuer H, Hegele M (2014) Age-related variations of visuo-motor adaptation beyond explicit knowledge. Front Aging Neurosci 6:1–12.

Heuninckx S, Wenderoth N, Debaere F, Peeters R, Swinnen SP (2005) Neural Basis of Aging?: The Penetration of Cognition into Action Control. 25:6787–6796.

Heuninckx S, Wenderoth N, Swinnen SP (2008) Systems neuroplasticity in the aging brain: recruiting additional neural resources for successful motor performance in elderly persons. J Neurosci 28:91–99.

Huang J, Gegenfurtner KR, Schütz AC (2017) Age effects on saccadic adaptation?: Evidence from different paradigms reveals specific vulnerabilities. 17:1–18.

Huang VS, Haith A, Mazzoni P, Krakauer JW (2011) Rethinking Motor Learning and Savings in Adaptation Paradigms: Model-Free Memory for Successful Actions Combines with Internal Models. Neuron 70:787–801.

Hulst T, van der Geest JN, Th??rling M, Goericke S, Frens MA, Timmann D, Donchin O (2015) Ageing shows a pattern of cerebellar degeneration analogous, but not equal, to that in patients suffering from cerebellar degenerative disease. Neuroimage 116:196–206.

Imamizu H, Tamada T, Yoshioka T, Pu B (2000) Human cerebellar activity re ¯ ecting an acquired internal model of a new tool. 403.

Izawa J, Shadmehr R (2011) Learning from Sensory and Reward Prediction Errors during Motor Adaptation. PLoS Comput Biol 7:e1002012.

Jonas E, Kording KP (2017) Could a Neuroscientist Understand a Microprocessor?? :1–24.

Kasuga S, Hirashima M, Nozaki D (2013) Simultaneous Processing of Information on Multiple Errors in Visuomotor Learning. 8.

Kim H, Morehead JR, Parvin DE, Moazzezi R, Ivry RB (2018) Invariant errors reveal limitations in motor correction rather than constraints on error sensitivity. Nat Commun Biol.

King BR, Fogel SM, Albouy G, Doyon J (2013) Neural correlates of the age-related changes in motor sequence learning and motor adaptation in older adults. Front Hum Neurosci 7:142.

Kording KP, Wolpert DM (2004) Bayesian integration in sensorimotor learning. Nature 427:244–247.

Körding KP, Wolpert DM (2004) Bayesian integration in sensorimotor learning. Nature 427:244–247.

Krakauer JW, Ghazanfar AA, Gomez-Marin A, Maciver MA, Poeppel D (2017) Neuron Perspective Neuroscience Needs Behavior: Correcting a Reductionist Bias. Neuron 93:480–490.

Krakauer JW, Ghez C, Ghilardi MF (2005) Adaptation to visuomotor transformations: consolidation, interference, and forgetting. J Neurosci 25:473–478.

Malone LA, Bastian AJ (2015) Age-related forgetting in locomotor adaptation. Neurobiol Learn Mem.

Marko MK, Haith AM, Harran MD, Shadmehr R (2012) Sensitivity to prediction error in reach adaptation. J Neurophysiol 108:1752–1763.

Mattay V, Fera F, Tessitore A, Hariri A, Das S, Callicott J, Weinberger D (2002) Neurophysiological correlates of age-related changes in human motor function. Neurology 58:630–635.

Mazzoni P, Krakauer JW (2006) An implicit plan overrides an explicit strategy during visuomotor adaptation. J Neurosci 26:3642–3645.

McDougle SD, Bond KM, Taylor JA (2015) Explicit and Implicit Processes Constitute the Fast and Slow Processes of Sensorimotor Learning. J Neurosci 35:9568–9579.

McDougle SD, Bond KM, Taylor XJA (2017) Implications of plan-based generalization in sensorimotor adaptation. :383–393.

McNab F, Klingberg T (2008) Prefrontal cortex and basal ganglia control access to working memory. Nat Neurosci 11:103–107.

Morehead JR, Qasim SE, Crossley MJ, Ivry R (2015a) Savings upon Re-Aiming in Visuomotor Adaptation. J Neurosci 20:14386–14396.

Morehead JR, Qasim SEE, Crossley MJJ, Ivry RB (2015b) Savings upon Re-Aiming in Visuomotor Adaptation. J Neurosci 35:14386–14396.

Morehead JR, Taylor JA, Parvin D, Ivry RB (2017) Characteristics of Implicit Sensorimotor Adaptation Revealed by Task-irrelevant Clamped Feedback. J Cogn Neurosci 26:194–198.

Morton SM, Bastian AJ (2006) Cerebellar Contributions to Locomotor Adaptations during Splitbelt Treadmill Walking. 26:9107–9116.

Nozaki D, Kurtzer IL, Scott SH (2006) Limited transfer of learning between unimanual and bimanual skills within the same limb. Nat Neurosci 9:1364–1366.

Nozaki D, Yokoi A, Kimura T, Hirashima M, Orban de Xivry J-J (2016) Tagging motor memories with transcranial direct current stimulation allows later artificially-controlled retrieval. Elife 5:e15378.

Oldfield RC (1971) The assessment and analysis of handedness: The Edinburgh inventory. Neuropsychologia 9:97–113.

Orban de Xivry J-J, Lefèvre P (2015) Formation of model-free motor memories during motor adaptation depends on perturbation schedule. J Neurophysiol 113:2733–2741.

Ostry DJ, Darainy M, Mattar AAG, Wong J, Gribble PL (2010) Somatosensory plasticity and motor learning. J Neurosci 30:5384–5393.

Pernet CR, Wilcox R, Rousselet GA (2013) Robust correlation analyses?: false positive and power validation using a new open source Matlab toolbox. 3:1–18.

Randolph C, Tierney MC, Mohr E, Chase TN (1998) The Repeatable Battery for the Assessment of Neuropsychological Status (RBANS): Preliminary Clinical Validity. J Clin Exp Neuropsychol 20:310–319.

Raz N, Lindenberger U, Rodrigue KM, Kennedy KM, Head D, Williamson A, Dahle C, Gerstorf D, Acker JD (2005) Regional brain changes in aging healthy adults: General trends, individual differences and modifiers. Cereb Cortex 15:1676–1689.

Schmahmann JD (2018) Neuroscience Letters The cerebellum and cognition. Neurosci Lett:0–1.

Seidler RD (2006) Differential effects of age on sequence learning and sensorimotor adaptation. Brain Res Bull 70:337–346.

Seidler RD (2007) Aging affects motor learning but not savings at transfer of learning. Learn Mem 14:17–21.

Shadmehr R, Holcomb H (1997) Neural correlates of motor memory consolidation. Science (80-) 277:821–825.

Shadmehr R, Krakauer JW (2008) A computational neuroanatomy for motor control. Exp Brain Res 185:359–381.

Shadmehr R, Mussa-Ivaldi F a (1994) Adaptive representation of dynamics during learning of a motor task. J Neurosci 14:3208–3224.

Shadmehr R, Smith M a, Krakauer JW (2010) Error correction, sensory prediction, and adaptation in motor control. Annu Rev Neurosci 33:89–108.

Shmuelof L, Huang VS, Haith AM, Delnicki RJ, Mazzoni P, Krakauer JW (2012) Overcoming Motor “Forgetting” Through Reinforcement Of Learned Actions. 32:14617–14621.

Sing G, Najafi B, Adewuyi A (2009) A novel mechanism for the spacing effect: Competitive inhibition between adaptive processes can explain the increase in motor skill retention associated with prolonged inter-trial spacing. Adv Mot Learn Mot Control:0–1.

Smith M a, Shadmehr R (2005) Intact ability to learn internal models of arm dynamics in Huntington’s disease but not cerebellar degeneration. J Neurophysiol 93:2809–2821.

Sombric CJ, Harker HM, Sparto PJ, Torres-oviedo G (2017) Explicit Action Switching Interferes with the Context-Specificity of Motor Memories in Older Adults. 9.

Sugiura M (2016) Functional neuroimaging of normal aging?: Declining brain, adapting brain. Ageing Res Rev 30:61–72.

Taylor J a., Krakauer JW, Ivry RB (2014) Explicit and Implicit Contributions to Learning in a Sensorimotor Adaptation Task. J Neurosci 34:3023–3032.

Taylor J a, Ivry RB (2014) Cerebellar and prefrontal cortex contributions to adaptation, strategies and reinforcement learning.

Taylor JA, Ivry RB (2011) Flexible Cognitive Strategies during Motor Learning. PLoS Comput Biol 7:e1001096.

Taylor JA, Klemfuss NM, Ivry RB (2010) An explicit strategy prevails when the cerebellum fails to compute movement errors. Cerebellum 9:580–586.

Taylor JA, Mcdougle SD (2016) Mental Rotation as a Behavioral and Neural Model of Explicit Aiming During Visuomotor Learning. :0–1.

Trewartha KM, Garcia A, Wolpert DM, Flanagan JR (2014) Fast But Fleeting: Adaptive Motor Learning Processes Associated with Aging and Cognitive Decline. J Neurosci 34:13411–13421.

Van Beers RJ, Wolpert DM, Haggard P (2002) When feeling is more important than seeing in sensorimotor adaptation. Curr Biol 12:834–837.

Vogel EK, Mccollough AW, Machizawa MG (2005) Neural measures reveal individual differences in controlling access to working memory. Nature 438:500–503.

Walhovd KB, Westlye LT, Amlien I, Espeseth T, Reinvang I, Raz N, Agartz I, Salat DH, Greve DN, Fischl B, Dale AM, Fjell AM (2011) Consistent neuroanatomical age-related volume differences across multiple samples. NBA 32:916–932.

Wei K, Kording KP (2009) Relevance of error: what drives motor adaptation? J Neurophysiol 101:655–664.

Wolpe N et al. (2016) Ageing increases reliance on sensorimotor prediction through structural and functional differences in frontostriatal circuits. Nat Commun 7:13034.

Wolpe N, Ingram JN, Tsvetanov KA, Henson RN (2018) Motor learning decline with age is related to differences in the explicit memory system Running title?: Motor learning with ageing. 44:1–39.

Wolpert DM, Miall RC, Kawato M (1998) Internal models in the cerebellum. 2:338–347.

Wong JD, Kistemaker DA, Chin A, Gribble PL (2012) Can proprioceptive training improve motor learning?? :3313–3321.

